# Three-dimensional motions of GroEL during substrate protein recognition

**DOI:** 10.1101/2022.09.15.508192

**Authors:** Kevin Stapleton, Tomohiro Mizobata, Naoyuki Miyazaki, Tomohiro Takatsuji, Takayuki Kato, Kenji Iwasaki, Daron M Standley, Takeshi Kawamura, Takanori Nakane, Junichi Takagi, Eiichi Mizohata

## Abstract

GroEL is a bacterial chaperonin responsible for the assisted folding of non-native and misfolded polypeptides into biologically active proteins. The adaptive nature of the recognition mechanism of chaperonins toward client polypeptides inherently lends itself to structural heterogeneity, which hampers detailed analyses of intermolecular recognition and binding. In this report, we used single-particle cryo-EM and multiple rounds of focused mask three-dimensional classification to reveal a landscape of distinct snapshots of endogenous GroEL complexed with an unfolded substrate, the water-soluble domain of human UDP glucuronosyltransferase 1A (UGT1A), at 2.7–3.5 Å resolution. We demonstrate that UGT1A occupies the GroEL ring asymmetrically, engaging with 2–3 contiguous subunits and that a subunit bound to UGT1A exhibits a wider range of conformational dynamics, consistent with AlphaFold models. These data reveal molecular motions during initial substrate capture at near-atomic detail.

## 1. Introduction

Chaperonin-assisted protein folding is essential for maintaining cellular homeostasis and protection against toxic protein aggregation. The chaperonin family contains bacterial GroEL and the structurally and functionally related eukaryotic protein Hsp60. The GroEL reaction cycle consists of alternating rounds of substrate recognition, nucleotide binding, encapsulation, and release of folded protein [1, 2]. The conformational changes that occur upon substrate capture, allowing recruitment of co-chaperonin GroES, facilitate the transfer of the substrate from a hydrophobic pocket to a hydrophilic chamber, wherein the substrate folds unencumbered by the cellular environment. Although numerous structural studies have revealed key conformational states of this process [2–6], most have been obtained in the absence of bound substrates, or where substrates were present, have been reported at resolutions that hinder detailed molecular interpretation of the reaction cycle.

Recent technological breakthroughs in cryogenic electron microscopy (cryo-EM) and innovations in single-particle analysis have enabled complex molecular machines to be studied at unprecedented resolution. In contrast to X-ray crystallography, a distinguishing feature of the single-particle approach is the ability to cluster particle images based on compositional or conformational heterogeneity. This feature, in combination with emerging computational methods and standardisation, has enabled the routine structural reconstruction of macromolecular assemblies at resolutions better than 4 Å.

In this report, we describe a complex between GroEL and substrate that formed endogenously in *E. coli* and was subsequently purified as a stable arrested complex. The substrate consists of the water-soluble domain of human UDP glucuronosyltransferase 1A (UGT1A), His6-tag, thioredoxin (TrxA) tag, and a TEV cleavage site. The resulting structures, ranging in resolution from 2.7–3.5 Å, constitute the most precise observations of a GroEL-substrate complex by cryo-EM.

We observed that UGT1A was asymmetrically bound within the GroEL ring, contacting 2 to 3 contiguous subunits. Focused mask classification segregated two distinct GroEL-substrate complexes: a 3.26 Å resolution GroEL-UGT1A complex corresponding to a state where UGT1A occupies a single ring of the binary complex and a 2.7 Å resolution complex corresponding to a state where UGT1A occupies both rings. Further application of focused mask classification on empty and occupied ring protomers (subunits) enabled the characterisation of additional distinct subunit conformational states. These observations indicate greater structural heterogeneity in the substrate-bound subunits compared with that in the unbound subunits. Interestingly, apical domain conformational changes correlated with subtle structural modifications in the nucleotide-binding pocket, suggesting that substrate and ATP binding are allosterically coupled. We used a hierarchical masking approach to subclassify particle images by altering the position of the focused mask during classification. This classification scheme enabled the isolation of new sub-clusters of particle images corresponding to subtle changes in GroEL composition and conformation and improved resolution of bound UGT1A, thus providing a detailed picture of chaperonin-substrate binding interactions. Overall, these data provide a dynamic 3D view of the chaperonin-assisted protein folding landscape.

## 2. Results

### 2.1 Purification of the GroEL-UGT1A complex

Initially, His_6_-TrxA-tagged UGT1A was overexpressed in *E. coli* (**Supplementary Fig. 1a**) and purified using nickel-affinity and size-exclusion chromatography. When the purified sample was incubated with TEV protease and then subjected to SDS-PAGE analysis, two clear bands at approximately 70 and 26 kDa were observed and corresponded to intact His_6_-TrxA-UGT1A monomer and TEV protease (**Supplementary Fig. 1b**). Thus, TEV did not cleave the fusion protein at the desired position. Blue native PAGE analysis revealed that the purified sample exhibits a large apparent molecular mass of 720 kDa over a wide range of pH values (pH 3.0−9.0) except for high alkalinity (pH 12.0), suggesting that the sample is in an oligomeric state (**Supplementary Fig. 1c**). Surprisingly, liquid chromatography-tandem mass spectrometry (LC-MS/MS) analysis revealed that the purified sample was a mixture of the human UGT1A and *Escherichia coli* chaperonin GroEL (**Supplementary Fig. 1d**). In subsequent purification attempts, this contaminating GroEL was extremely difficult to remove, which prompted us to hypothesize that His_6_-TrxA-UGT1A and GroEL may stably associate within the bacterial cytosol. Negative-stain EM was used to validate the quality of the GroEL-UGT1A complex, resulting in discrete particulates in multiple orientations evenly distributed in the field of view (**Supplementary Fig. 1e**). Furthermore, there was visible density within the GroEL ring, suggesting bound UGT1A (**Supplementary Fig. 1e**).

**Fig. 1.**
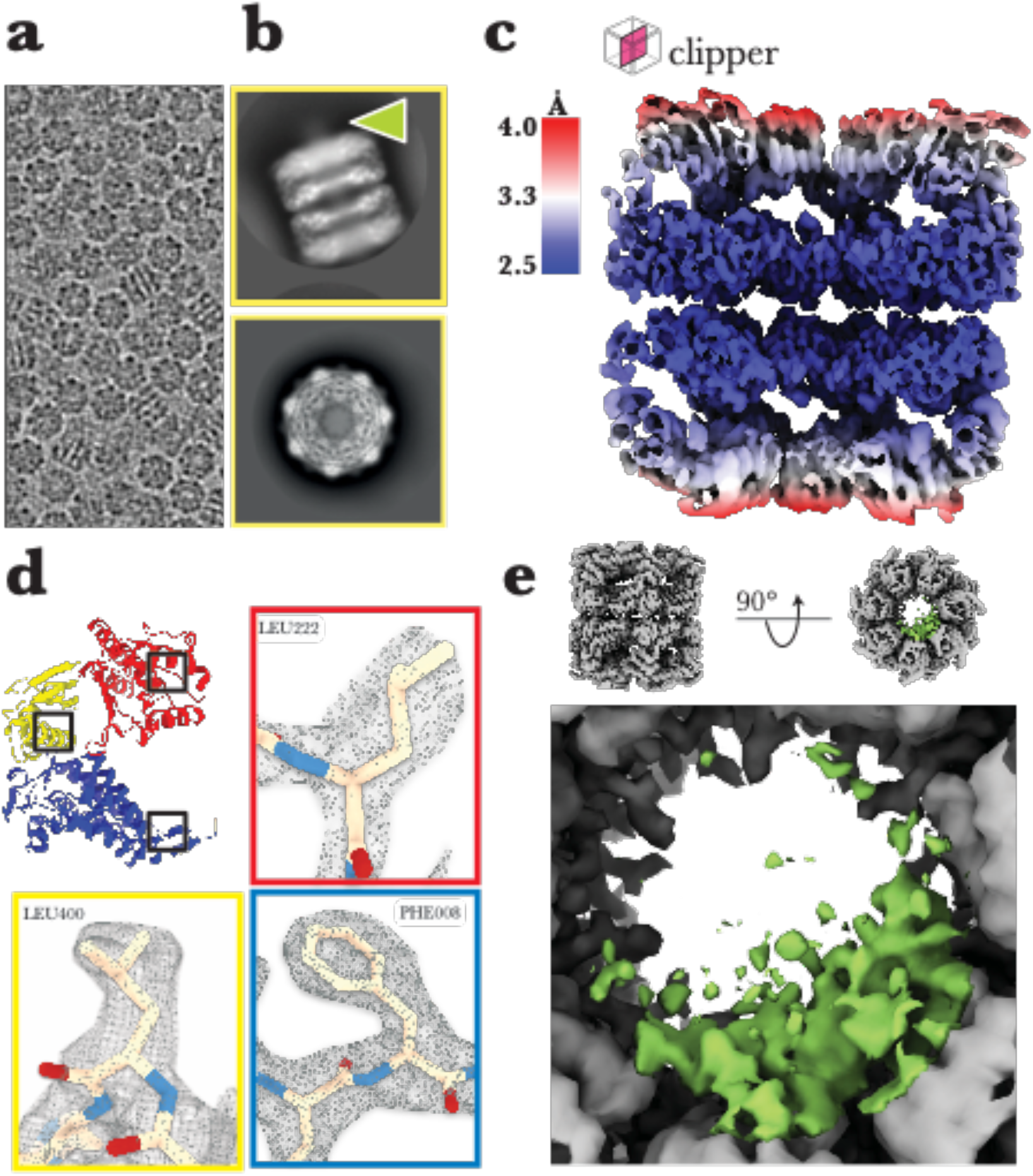
The 2.8 Å consensus refinement. Asymmetric reconstruction and atomistic model of the GroEL/UGT1A complex. **a**, Typical motion-corrected and CTF estimated raw micrograph containing visible GroEL particles suspended in multiple orientations alongside reference-free 2D class averages (2/38) from extracted particle images **b**, Side-on and top views of GroEL/UGT1A with visible ring occupation by flexible non-native UGT1A in the side-on view (green arrow). **c**, The 2.8 Å resolution cryo-EM density map is coloured by local resolution in the clipper view. **d**, A single GroEL modelled subunit coloured according to the domains: apical (red), intermediate (yellow), and equatorial (blue), with side-chain fitting of the EM density map (grey mesh) of three residues in the apical (Ala223), intermediate (Leu400), and equatorial domains (Phe008). **e**, Bottom-end view of the GroEL-UGT1A consensus map (dark grey). The substrate UGT1A (green) density forms asymmetrically to one side of the ring cavity.

### 2.2 Overall structure of the GroEL-UGT1A complex

The sample containing the GroEL-UGT1A complex was prepared for cryo-EM imaging and analysed as described in Methods and Supplementary Materials. From 2,951 recorded movies, individual particles were observed to be uniformly distributed and in multiple orientations (**Fig. 1a**) in the field of view. After motion-correction, the contrast transfer function (CTF) parameters of each micrograph were estimated [7, 8] and processed in RELION 3.1 [9]. 1,202,096 particle images were successfully extracted and classified into 50 reference-free two-dimensional (2D) classes displaying multiple particle orientations with flexible substrate UGT1A density identifiable in the ring cavity from side-on and top-view 2D projections (**Fig. 1b**).

Structurally, each GroEL protomer was divided into three major domains: (i) the equatorial domain, which forms the interface between the two rings and contains a site for nucleotide binding and hydrolysis; (ii) the apical domain, which lines the ring cavity and is the site for binding of co-chaperonin GroES and initial non-native substrate capture; and (iii) the intermediate domain, which acts as a hinge connecting the equatorial and apical domains and facilitates extensive nucleotide- and ligand-induced conformational rearrangement of the subunits required for a typical folding reaction cycle. The GroEL complex adopts D7 symmetry because the ring comprises seven identical subunits. However, because previous studies have suggested that substrate binding can occur asymmetrically and that each subunit within a heptameric ring can move independently from other subunits [4, 10], we conducted all subsequent analyses without imposing symmetry on particle images. Thus, we deliberately allowed conformational or compositional heterogeneity in particle images to be retained for subsequent separation by focused mask three-dimensional (3D) classification (focus classification). This approach resulted in a GroEL-UGT1A consensus refinement at an estimated global resolution of 2.8 Å by Fourier shell correlation (FSC) with a cut-off at 0.143 [11–14] (**Supplementary Fig. 3**).

Local resolution distribution throughout the refined 2.8 Å consensus map (**Fig. 1c**) revealed a gradual decrease from the equatorial domains (2.5 Å) to the apical domains (3.4 Å), indicating heterogeneity and/or flexibility in the latter, consistent with previous reports of continuous domain-level motions within the GroEL subunit [10, 15, 16].

Despite the modest resolution, the map quality was consistent throughout the structure, and most secondary structure elements were unambiguously resolved in all three major domains. Furthermore, distinct side-chain densities were visible in many instances (**Fig. 1d**), enabling model building with an unambiguous residue register. In addition, UGT1A-associated densities were observed inside the GroEL ring (**Fig. 1e**). Interestingly, the UGT1A density was markedly more present in one ring (denoted “bottom”) compared with that in the other (denoted “top”; **Supplementary Fig. 4**) and was asymmetrically distributed within the bottom ring, contacting 2–3 neighbouring apical domain subunits (**Fig. 1e**).

### 2.3 Observation of single- and double-occupied rings

We used focused mask 3D classification using polyshape masks to investigate further the asymmetric distribution of UGT1A in the 2.8 Å consensus reconstruction (**Fig. 1e**; see **Methods** for details). This classification process gave two discrete 3D volume classes: a dominant 3D volume containing ∼1.1 M (96%) of all particle images and an unexpected minority class containing ∼22,000 (2%) particle images (**Fig. 2a**). In the dominant class, the distribution of UGT1A density in the ring cavity was more pronounced and engaged with 2–3 contiguous subunits, whereas UGT1A density was predominantly absent for apical domains located on the opposing side of the cavity (**Supplementary Fig. 5**). This asymmetric distribution strongly supports a model where substrate binding occurs between 2–3 apical domains within a heptameric ring and UGT1A.

**Fig. 2.**
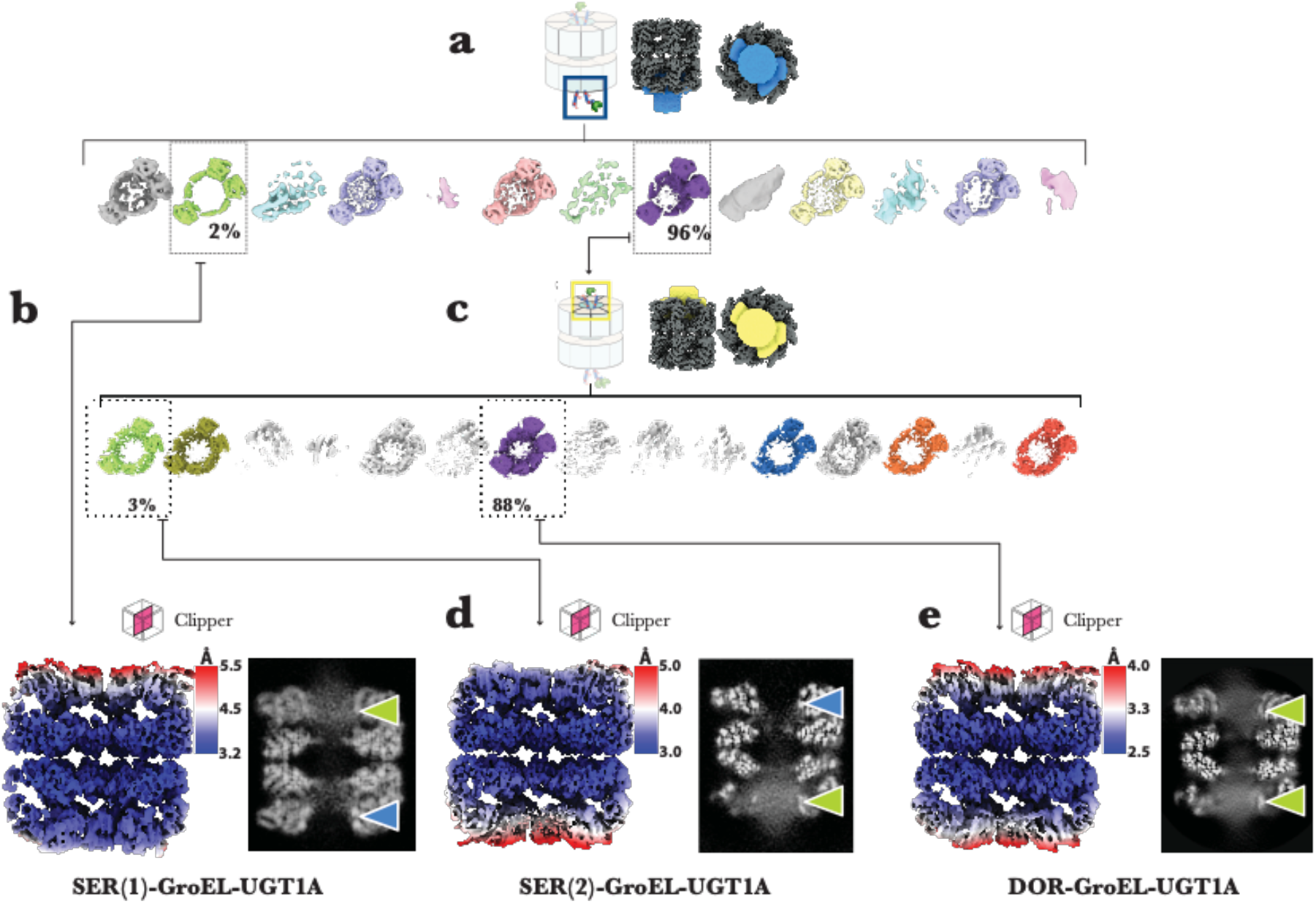
Separation by focus classification of isolated GroEL-UGT1A complexes with UGT1A bound to one or both rings. **a**, The GroEL-UGT1A consensus particles were used in the first round of focus classification. The bottom ring of the GroEL-UGT1A 3D density volume is covered with a soft mask, yielding two discrete 3D volume particle classes (boxed): an occupied density volume with 96% of particles (purple) and an empty density volume with 2% of particles from the consensus map (green). **b**, Reconstruction and refinement of the SER(1)-GroEL-UGT1A complex from the empty volume in the first round of focus mask classification. The 3D volume is coloured by local resolution and shown in clipper view and 2D z-slice. By removing compositional heterogeneity in the bottom ring, the local resolution distribution is noticeably more uniform compared with that in the top ring. A 2D z-slice through the experimental images is shown to the right, and flexible UGT1A occupies the top ring (green arrow) while the bottom ring remains empty (blue arrow). **c**, The second round of focus classification is performed on the reconstructed volume containing 96% of the particle images. A soft mask is placed on the top ring, yielding two discrete 3D volume particle classes (boxed): an occupied density volume with 88% of particles (purple) and an empty density volume with 3% of particle images (green). **d**, Reconstruction and refinement of the SER(2)-GroEL-UGT1A complex from the empty volume in the second round of focus mask classification. The 3D volume is coloured by local resolution and shown in clipper view and 2D z-slice. As was visible in (**c**), the local resolution distribution is noticeably more uniform in the area ‘cleaned’ of compositional heterogeneity, i.e., the local resolution in the top ring is more uniform than in the bottom ring. A 2D z-slice through the experimental images is shown to the right, and flexible UGT1A occupies the bottom ring (green arrow) while the top ring remains empty (blue arrow). **e**, Reconstruction and refinement of the DOR-GroEL-UGT1A complex from the occupied volume in the second round of focus mask classification. The 3D volume is coloured by local resolution and shown in clipper view and 2D z-slice. By purging the reconstruction of SER complexes, the local resolution distribution for the rings of this complex indicates conformational heterogeneity caused by domain-level flexibility in the subunits rather than compositional variation. A 2D z-slice through the experimental images is shown on the right, and flexible UGT1A occupies both rings (green arrows).

In addition to the dominant class, a minority class (∼2% of particle images) was observed wherein no observable UGT1A density was present in the ring. To further explore this unexpected result, we removed the focused mask from the minority class and refined the associated particle images without angular reassignment (**Methods**) (**Fig. 2b**). This resulted in a 3.5 Å resolution GroEL binary ring complex where one ring was populated with visible UGT1A density, and the other was devoid of any UGT1A density. 3D visualisation of the volume and 2D/3D slices (**Fig. 2b**) of this density map confirmed the isolation of a subpopulation of the Single-Empty-Ring (SER) GroEL-UGT1A structure complex (designated SER(1)-GroEL-UGT1A) from the consensus map particle image population. We immediately expanded this strategy to the dominant class and performed another round of focus classification (**Fig. 2c**) with the focused mask positioned over the top ring.

The second round of focus classification resulted in two additional discrete 3D volumes (**Fig. 2c**): a minor 3D class volume containing a subpopulation of ∼36,000 empty-ring particle images (3%) and a dominant 3D volume class containing a subpopulation of ∼1 M occupied-ring particle images (88%). Removal of the mask and separate refinements of these two particle classes identified a second SER(2)-GroEL-UGT1A complex at 3.26 Å global resolution (**Fig. 2d**) and a Double-Occupied-Ring (DOR) GroEL-UGT1A complex with an improved global resolution of 2.7 Å (**Fig. 2e**). Visualisation of the refined 3D volumes and inspection of particle images through z-slices confirmed the single-/double-ring occupation. These structural complexes represent a 17:1 DOR:SER image population. Interestingly, when we superimposed empty rings over occupied rings from all refinements, a noticeable elevation of the substrate-binding helices (H and I) lining the occupied ring cavity was observed.

### 2.4 Substrate binding causes apical domain elevation

To further investigate conformational changes in the occupied apical domains, we generated a soft mask around a single protomer from both the structure of the UGT1A empty-ring of the SER(2)-GroEL-UGT1A complex (**Fig. 3b**) and the UGT1A occupied-ring of the DOR-GroEL-UGT1A complex (**Fig. 3c**). A single round of focus classification (**Fig. 3c**) without particle alignment on the occupied-ring (OR) protomer from DOR-GroEL-UGT1A yielded four discrete conformations designated OR1, OR2, OR3, and OR4 (**Fig. 3e**). These four conformers represented snapshots within a continuous range of motion. Assigning atomic coordinates to the focused maps (**Fig. 3g**) revealed an 8.5–9.2 Å elevation in the substrate and ligand binding helices H and I. In contrast, focus classification of the SER(2)-GroEL-UGT1A Empty-Ring (ER) protomer gave a single discrete conformation (**Supplementary Fig. 6**). However, the limited number of available particle images contained in the 3.26 Å SER(2)-GroEL-UGT1A reconstruction (C1) may have masked the possible conformational variation, as previously reported for apo-GroEL [10].

**Fig. 3.**
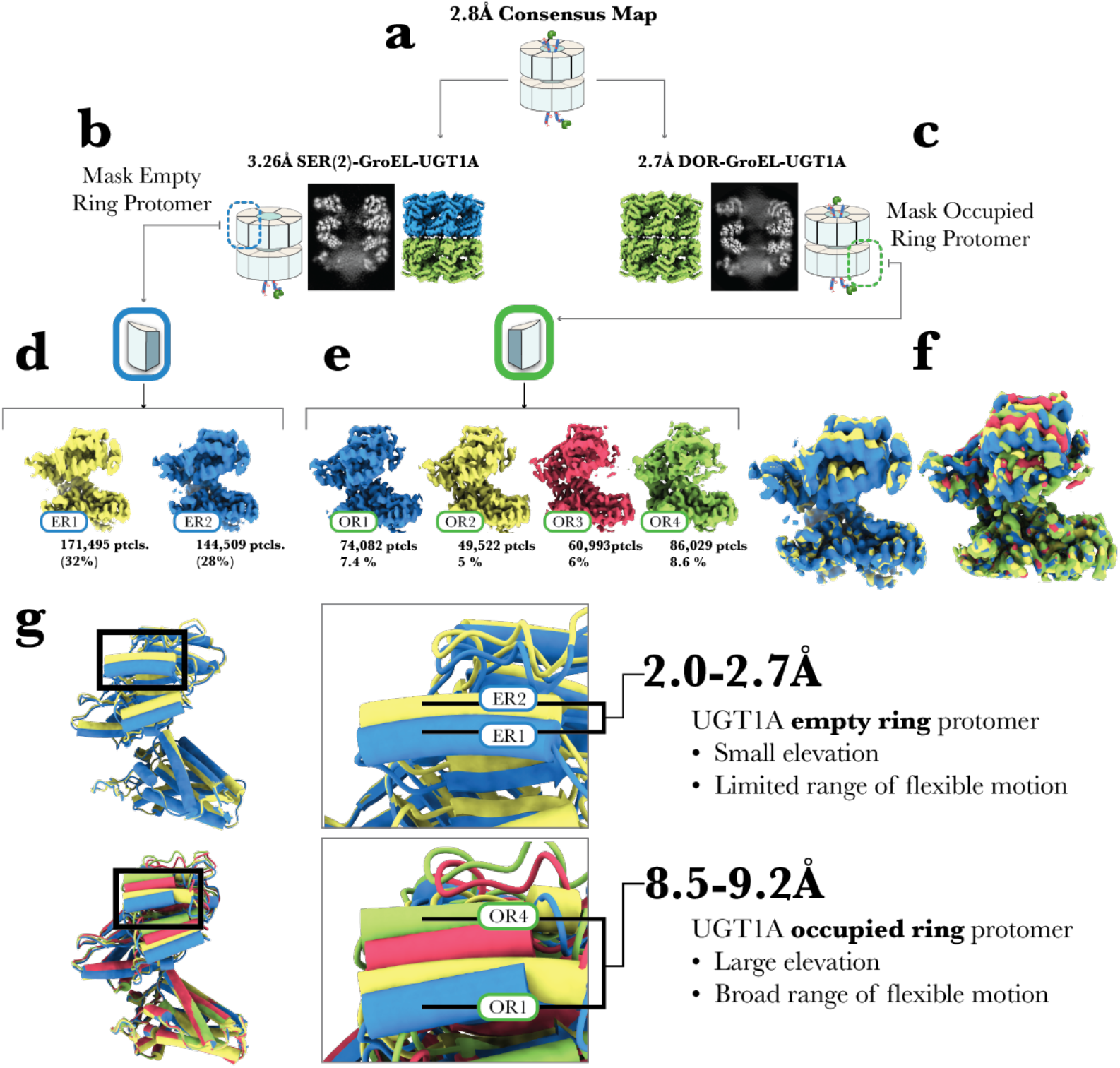
En bloc apical domain elevation. **a**, The 2.8 Å consensus map before separating substrate empty and occupied ring GroEL-UGT1A complexes by focus classification. **b**, The 3.26 Å resolution SER(2)-GroEL-UGT1A complex is accompanied by the 2D z-slice of the 3D volume. The complex is coloured by empty (blue) or occupied (green) ring states. A single protomer from the empty ring is masked for focus classification in the SER(2) complex (left, blue dashed square). **c**, The 2.7 Å resolution DOR-GroEL-UGT1A complex is accompanied by the 2D z-slice of the 3D volume. The complex is coloured by the substrate-occupied ring (green). A single protomer from the occupied ring is masked for focus classification in the DOR complex (left, green dashed square). **d**, Focus classification of the empty ring protomer yields two discrete 3D volume classes (ER1 and ER2) containing 32% and 28% of the SER(2)-GroEL-UGT1A particle image population. **e**, Focus classification of the occupied ring protomer yields four discrete 3D volume classes (OR1–4) containing 27% of the DOR-GroEL-UGT1A particle image population. **f**, Superimposition of the 3D density volumes from both classifications. The elevation in the apical domains is visible when the 3D density volumes from both classes are superimposed, although it is more pronounced in the OR protomer than in the ER protomer. **g**, The fitted and refined atomic coordinates are displayed in cylinders and stubs from the ER and OR density volumes. En block apical domain elevation is exhibited most prominently in the substrate-binding helices H and I. OR protomers display a broader range of continuous elevation (8.5–9.2 Å across residues in helix H) than ER protomers (2.0–2.7 Å across residues in helix H), as seen in the enlarged pictures of helix H from OR and ER models.

This issue was addressed by aligning SER(2)-GroEL-UGT1A images along the D7 symmetrical axis, refining the map imposing D7 symmetry, and expanding the resulting particle projections (**Supplementary Fig. 7**). We then generated a new soft mask around a single protomer and performed another round of focused mask classification over a protomer from the SER(2)-GroEL-UGT1A (D7) reconstruction. As shown in **Fig. 3b,d**, this analysis afforded two discrete conformers (ER1 and ER2). After fitting the atomic coordinates to these volumes, an elevation of 2.0–2.7 Å (**Fig. 3g**) for the substrate-binding helices was observed (**Movie 1**).

### 2.5 Visualization of bound UGT1A

Removal of the soft mask from OR1–4 focused maps (see **Methods**) and separate refinement of these image stacks without particle alignment (retaining all the original Euler angles assigned to each particle image from the DOR complex) resulted in four new DOR-GroEL-UGT1A complexes ranging between 3.2–3.3 Å resolution (**Supplementary Fig. 8**), referred to as OR(1–4)-GroEL-UGT1A, reflecting classes (OR1–4) and their conformational changes in the context of the GroEL molecule from which they were derived.

The resolution of bound UGT1A, previously obstructed by the focus mask, was noticeably improved and individual protomers were observed to protrude from the GroEL ring in a stepwise manner breaking the ring symmetry and leaving neighbouring subunits unaffected by the *en bloc* apical domain elevation (**Supplementary Fig. 9**). From the OR4-GroEL-UGT1A complex structure reported at 3.2 Å resolution (global), UGT1A was unambiguously observed binding to α-helix I (res: 256–259) and contacting the US-loop (res: 202–204) (**Fig. 4, Movie 3**).

**Fig. 4.**
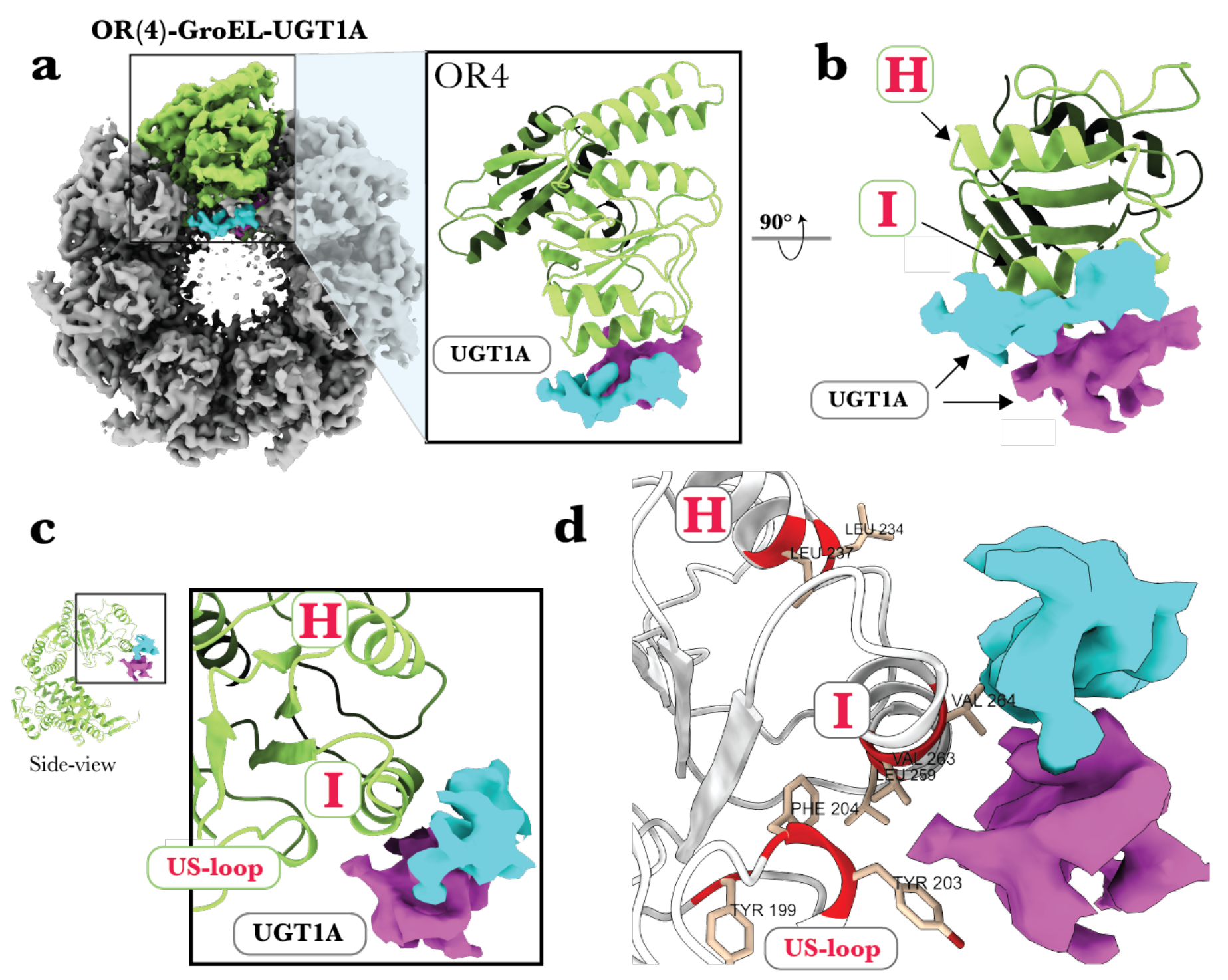
UGT1A binding density to a GroEL ring protomer (subunit). **a**, The OR(4)-GroEL-UGT1A complex density volume (grey), with non-continuous UGT1A density (cyan and purple) seen in the GroEL ring contacting the OR4 protomer. The position of the UGT1A substrate density in relation to the OR4 fitted atomic coordinates (modelled in ribbon) is displayed on the right (green). **b**, The substrate-binding helices H and I are now front-facing after rotating the OR protomer by 90°. The density of UGT1A forms primarily at and below helix I. **c**, Side-on view of the OR4 protomer with the magnified apical domain where UGT1A forms (inset). The density of UGT1A is below helix I and the US-loop. **d**, OR protomer (white) and essential hydrophobic substrate-binding residues (red).

## 3. Discussion

Previous structural investigations of GroEL-substrate complexes [6, 16–22] have generally corroborated biochemical reports [23, 24], which hypothesize that 2–3 contiguous subunits are required for substrate interaction [4, 6, 25–27]. Among these reported structures, all except two [21, 27] were performed by rapid mixing as opposed to endogenous capture. Two previous reports of structures with resolution better than 4 Å [16, 28] used model peptides as surrogate substrates. The question of how GroEL recognizes substrates *in cellulo* has thus remained open. In this study, we presented a single-particle cryo-EM reconstruction of endogenous chaperonin GroEL spontaneously complexed with non-native substrate UGT1A at near-atomic resolution.

In our consensus reconstruction, occupation of the UGT1A substrate appeared to contact 2–3 contiguous subunits in a ring (**Fig. 1e**). This asymmetry was further confirmed by systematically separating UGT1A empty and occupied ring complexes by a focused mask classification strategy (**Fig. 2, Methods**, and **Supplementary Fig. 9**).

The final ‘focus-cleaned’ maps SER(1/2)-GroEL-UGT1A (3.26 Å/3.5 Å) and DOR-GroEL-UGT1A (2.7 Å) all exhibited an asymmetric occupation of UGT1A in their respective rings. Interestingly, the top-ring of the DOR-GroEL-UGT1A complex, although occupied with the substrate, was uniformly distributed around the ring with no discernible pattern of occupation. The observation that one ring has a clear asymmetric binding motif while the other does not suggests that, despite the highly flexible nature of unfolded UGT1A and the more prominent and overpowering signal of the GroEL complex, alignment of particle images during refinement occurred with respect to structural features in the bottom ring and not the top ring.

Detection of non-native substrate protein complexed with GroEL has been scrutinized by various methods [4, 17–19, 21, 29]. Of these investigations, there have been notable examples of substrate bound to one ring [4, 6, 21, 25, 27] and reports of variable substrate occupancy [22, 30]. Except for one report [26], structural isolation of chaperonin complexes with different substrate binding forms has remained limited. This limitation is caused by the symmetry restraints applied during analysis or moderate resolutions impeding the elucidation of discrete conformational states. The present study shows the potential of hierarchical masking to elucidate discrete states in a molecular system that exhibits compositional heterogeneity, conformational heterogeneity, or both.

Superimposition of empty ring density volumes over occupied ring density volumes (from all independently derived SER and DOR complexes) indicated an apparent ∼5 Å ring cavity expansion, in agreement with previous reports [25]. However, after atomic fitting and superimposition of atomic coordinates, the structural heterogeneity was attributed not to an expansion per se but to the elevation of the substrate-binding helices H and I (**Supplementary Fig. 10**), which was further supported by molecular modelling using AlphaFold (**Supplementary Fig. 14**). This revised model is also consistent with previous reports of the ring ‘narrowing’ in GroEL-RuBisCO and GroEL-glutamine synthetase complexes [6, 25].

A notable observation was the loss of density corresponding to the inter-subunit contact involving Arg231 (**Supplementary Fig. 10b**) between neighbouring subunits from a UGT1A-empty ring. Previously, a similar feature was observed in a 4 Å crystal structure [16]. In that investigation, a 12-mer peptide was bound primarily to a single subunit situated in the interhelical groove between H and I. This central interaction included contact with two arginines (Arg231 and Arg268) from neighbouring subunits on either side. Although molecular replacement of the Arg231 side-chain from the UGT1A-occupied ring was not unambiguous, we hypothesize that the substrate load within a ring cavity may break this Arg231 link by coordinating the substrate with 2–3 subunits, as seen previously [31], allowing the movements we observed in occupied but not empty rings (**Fig. 3g** and **Supplementary Movie 1**).

During a typical reaction cycle, GroEL heptameric ring protomers (subunits) exhibit intra- and inter-domain flexibility characterized by elevation and twisting in an ATP-dependent manner required for chaperonin function [32, 33]. Several detailed structural investigations have illustrated these large *en bloc* conformational rearrangements at various stages of the GroEL functional cycle [2, 5, 34]. However, initial substrate recognition and substrate-initiated conformational changes have only begun to be addressed, albeit with intriguing results that indicate apical domain rearrangements around a captured non-native substrate [26].

Despite the paucity of high-resolution structures of GroEL-substrate complexes, biochemical reports have demonstrated consistently that GroEL protomers contain significant flexibility, plasticity, asymmetry, and structural variation in the apical domains [15, 31, 35–37]. However, with the addition of a recent study by Roh et al., which presented a detailed investigation into the subunit motions from apo-GroEL by single-particle cryo-EM, the results described in this study bridge prior work by revealing both substrate empty and occupied GroEL rings from a single sample.

This analysis revealed a significantly larger degree of continuous elevation of GroEL ring cavities occupied with UGT1A (OR1–4) when compared with that of ring cavities that were without (empty) UGT1A (ER1–2) (**Fig. 3**). Moreover, ER1, ER2 and the subunits from Roh et al. displayed the same level of apical domain elevation, indicating that the restricted apical domain elevation is a feature of unoccupied empty ring motions. Additionally, although OR1 and OR2 contain substrate density, they superimpose with low RMSD over ER1 and ER2. This observation suggests that substrate binding to a subunit does not restrict subunit flexibility but is responsible for the increased elevation in OR3 and OR4 conformations. This finding was further supported by superimposing the unmasked maps of OR1–4 to reveal the stable, well superimposed, and unelevated neighbouring subunits to either side of the elevated subunits (**Supplementary Fig. 9b**). In addition to elevation in the apical domain, OR1–4 protomers exhibited ‘breathing’ motions in the nucleotide-binding pocket (**Supplementary Movie 2**), suggesting that the increased elevation exclusively seen in OR protomers allosterically modifies the ATP binding pocket (**Movie 2**) in the equatorial domain, despite the physical separation between the two.

We examined the conformational states of ATP-binding residues in the ER1–2 and OR1–4 subunits because downstream ATP-induced conformational changes are a prerequisite for GroES encapsulation of a substrate-occupied GroEL ring (**Supplementary Fig. 11a**). The nucleotide-binding pocket in the OR1–4 protomers was found to exhibit more structural changes than that of the ER1–2 protomers. Additionally, the focus map of OR4 contained a pronounced density near the expected position of the Asp87-coordinated Mg^2+^ in the occupied state but not in the empty protomer (**Supplementary Fig. 11b**). These subtle structural differences prompted us to investigate whether the predicted ATP binding geometry was affected by these changes.

The binding energies of ATP bound to individual subunits (**Supplementary Fig. 11c**) were computed by flexible docking of ATP to both the occupied and empty states using Glide software (see **Methods**). The best ATP docking energy for the ER protomers was –4.5 kcal/mol. In contrast, the best ATP docking energy for the OR protomers was –7.0 kcal/mol (**Supplementary Fig. 11d**). Even when ATP poses were filtered by a ligand RMSD ≤ 4.5 Å from the ATP-bound GroEL (PDB ID: 3WVL), the observed trend was the same: ATP binding energies in substrate-bound subunits were significantly lower (more stable) than those in unbound subunits. Moreover, the ATP binding energies appeared to increase (less stable) with the degree of domain elevation. We hypothesize that ATP binding affinity within a GroEL ring is highest when needed — that is, when the ring cavity is occupied but has not yet undergone an ATP-induced conformational change. The allosteric relationship between substrate and nucleotide binding is an important topic for further exploration.

Previous EM studies visualising exposed GroEL rings with a bound substrate have been reported as early as 1993 [17] and demonstrate a wide range of resolutions [4, 6, 21, 25–27]. Except for the latest report [38], these structures have not achieved resolutions sufficient to observe clear chaperonin-substrate interactions. This lack of resolution is caused by the mobility of an unfolded flexible protein, which averages out or misaligns during consensus refinement, thus inhibiting the reconstruction of high-resolution features from the heterogeneous area. Remarkably, from our discrete class of particles within the OR4 focused map (or4-GroEL-UGT1A, EMDB ID: EMD-33352), UGT1A density is observable just under helix I (residues 256–268), with additional density forming in front of the US-loop (**Fig. 4, Movie 3**). Although the EM reconstructions of GroEL binding helices presented here are too weak to visualize side-chain residues, UGT1A backbone density at the hydrophobic helix I and US-loop agrees with previous observations of a complex between GroEL and bound non-native malate dehydrogenase [4].

Moreover, when viewing the cavity region of OR4-GroEL-UGT1A, the side-chains Val264 and Tyr203 from helix I (**Fig. 4**) and the US-loop of GroEL face the UGT1A density and have been experimentally proven to be essential to substrate binding, folding, and ES encapsulation [24]. Additionally, Tyr203 is conserved in 49/50 HSP60 homologues. The UGT1A density was too ambiguous to model atomic coordinates. Nonetheless, based on indistinct but continuous density situated around helix I, a poly-alanine peptide consisting of 26 residues was placed into the substrate-associated density (PDB ID: 7XOM).

Lastly, reconstructing the OR focus maps without the masks yielded a series of new GroEL-substrate complexes. In all maps, we observed UGT1A bound to 2–3 subunits of the heptameric ring asymmetrically (**Supplementary Fig. 9c**). These maps also show an assembly or ‘load’ of UGT1A density localising prominently at a single subunit. Neighbours to either side of the central UGT1A bound subunit are not affected by the significant conformational changes that can be seen in the OR1–4 masked and unmasked maps (**Supplementary Fig. 9b**). This is an intriguing structural observation of substrate-bound GroEL complexes because others have reported subunit rearrangement upon polypeptide binding. However, what is observed in these reconstructions suggests that neither the substrate load nor the 9.3 Å elevation in the apical domain affects the conformation or arrangement of neighbouring subunits, resulting in only the substrate-bound subunit breaking the symmetry of the GroEL ring. The strong binding of UGT1A to a single GroEL subunit observed in OR4 may have disrupted the folding cycle, leading to the formation of this arrested complex.

The initial capture of non-native polypeptide occurs at the substrate-binding helices in the apical domains lining the open cavity. Upon substrate binding, ATP-dependent conformational changes facilitate association with the co-chaperonin GroES, which encapsulates a substrate-containing GroEL ring, and the folding reaction proceeds. However, low-resolution details have severely hampered studies describing the initial substrate capture and binding motifs.

With the tools uniquely embedded within the single-particle method, this study brings a wide range of previous observations into a single coherent picture while raising new questions about the role of substrate binding and initialisation of the GroE-assisted protein folding pathway. We reported the high-resolution refinement of multiple spontaneously formed endogenous GroEL-UGT1A complexes ranging from 2.7–3.5 Å resolution. We demonstrated that UGT1A binding is asymmetric within a GroEL ring-cavity, and UGT1A unambiguously contacts 2–3 contiguous subunits, strongly interacting above and below substrate-binding helix I of a central subunit’s apical domain. Moreover, we successfully observed hitherto undescribed, novel subunit motions in the apical domain and ATP binding pocket, uniquely exhibited by occupied ring protomers. Lastly, this study represents the first directed separation of high-resolution GroEL-substrate complexes (or GroEL complexes of any kind) by single-particle cryo-EM. When applied to previous and future GroEL data sets, this focus-cleaning strategy should further remove significant heterogeneity and improve resolvability and map quality, revealing hitherto unseen, biologically relevant GroEL features and/or structures [2, 4, 6, 38, 39].

## 4. Methods

### 4.1 GroEL-UGT1A expression and purification

This study was originally initiated to produce and solve the 3D structure of UGT1A. A cDNA fragment encoding the human UGT1A (Uniprot entry: P19224) soluble domain without the N-terminal signal sequence and the C-terminal membrane anchor was cloned into the expression vector pTAT6 (generously gifted by Dr Marco Hyvonen, University of Cambridge, UK). The resulting plasmid expresses N-terminal His_6_-TrxA-tagged UGT1A with a linker containing the tobacco etch virus (TEV) protease recognition site. The plasmid was co-transformed with the plasmid pKY206 carrying the GroEL/ES gene [40] into *Escherichia coli* strain BL21Star(DE3), and transformants were screened on Luria-Bertani (LB) agar supplemented with 50 μg/ml ampicillin and 20 μg/ml tetracycline. Cells were grown at 37 °C in an LB broth supplemented with 50 μg/ml ampicillin to an OD_600_ of 0.6, and protein expression was induced at 15 °C for 24 h by adding isopropyl-β-D-thiogalactopyranoside to a final concentration of 0.4 mM. The cells were harvested by centrifugation, resuspended in buffer A (phosphate-buffered saline (PBS), 0.5 M NaCl, 10% (w/v) sucrose, 10% (v/v) glycerol, 0.2 M L(+)-arginine hydrochloride, pH 7.5) and disrupted with an EmulsiFlex-C3 (Avestin). The crude extract was centrifuged at 100,000 × *g* for 30 min, and the supernatant was subjected to affinity purification with a His-Trap HP 5 mL (GE Healthcare) column. The His_6_-tagged protein was eluted with a 0–0.5 M imidazole gradient in buffer A. The eluted sample was concentrated and applied to a Superdex 200 16/60 (GE Healthcare) column equilibrated with buffer B (20 mM Tris-HCl, 0.15 M NaCl, pH 7.5). The purified protein was concentrated to 33 mg/ml. The amount of the protein was determined spectrophotometrically using *A*_280_ × 5.0 mg/ml. The purification procedure was performed at 4 °C.

Purified protein aliquots were incubated with or without TEV protease at 4 °C for a week and then subjected to sodium dodecyl sulfate-polyacrylamide gel electrophoresis (SDS-PAGE). The purified samples were also incubated in seven different buffers ranging in pH from 3.0 to 12.0 and then subjected to Blue Native PAGE using a NativePAGE™ Sample Prep Kit and a 3–12% Bis-Tris Gel (Invitrogen).

### 4.2 Liquid chromatography-tandem mass spectrometry

The purified protein was confirmed to be the GroEL-UGT1A complex by liquid chromatography-tandem mass spectrometry (LC-MS/MS) as follows. The samples were suspended in 25 vol% CH_3_CN/25 mM NH_4_HCO_3_, reduced in 1.2 mM Tris (2-carboxyethyl) phosphine for 15 min at 50 °C, and alkylated in 3 mM iodoacetamide for 30 min at room temperature. The samples were digested with 100 ng trypsin (Promega) overnight at 37 °C. After drying with a SpeedVac (Thermo Fisher Scientific), the peptides were dissolved in 0.2% CF_3_COOH/2% CH_3_CN. Tandem MS analysis was performed with an LTQ Orbitrap ELITE ETD mass spectrometer (Thermo Fisher Scientific). Aliquots of the trypsinized samples were injected onto a micro-precolumn, C18 PepMap 100 Peptide Trap cartridge (5 × 0.3 mm i.d., Thermo Fisher Scientific) attached to an injector valve for desalting and concentrating the peptides. After washing the trap with 98% H_2_O/2% CH_3_CN/0.2% CF_3_COOH, the peptides were loaded onto a separation capillary reverse phase column (L-column2 micro C18 column 3 μm, 200 Å, 150 × 0.2 mm i.d., CERI, Tokyo, Japan) by switching the valve. The eluents used were A, 98% H_2_O/2% CH_3_CN/0.1% HCOOH, and B, 10% H_2_O/90% CH_3_CN/0.1% HCOOH. The column was developed at a flow rate of 1.0 μL/min, with a concentration gradient of CH_3_CN: from 5% B to 35% B for 100 min, from 35% B to 95% B for 1 min, sustained at 95% B for 9 min, from 95% B to 5% B for 1 min, and finally re-equilibrated with 5% B for 9 min. Effluents were introduced into the mass spectrometer via a nanoelectrospray ion interface, which held the separation column outlet directly connected to a nanoelectrospray ionization needle (PicoTip FS360-50-30, New Objective Inc.). The electrospray ionization voltage was 2.0 kV, and the transfer capillary of the LTQ inlet was heated to 200 °C. The MS was operated in a data-dependent acquisition mode, in which the MS acquisition with a mass range of *m/z* 420–1,600 was automatically switched to MS/MS acquisition under the automated control of Xcalibur software. The top four precursor ions were selected by an MS scan with Orbitrap at a resolution of 240,000 for subsequent MS/MS scans by ion trap in the normal/centroid mode using the automated gain control mode with automated gain control values of 1 × 10^6^ and 1.00 × 10^4^ for full MS and MS/MS, respectively. We also employed a dynamic exclusion capability that allows the sequential acquisition of the MS/MS of abundant ions in the order of their intensities with an exclusion duration of 2.0 min and exclusion mass widths of –5 and +5 ppm. The trapping time was 100 ms with auto gain control on. Tandem mass spectra were extracted using the Proteome Discoverer version 2.1. All MS/MS samples were analysed using Mascot (Matrix Science, version 2.6). Mascot was set up to search the SwissProt_2022_01 database (selected for *Homo sapiens*, 20,377 entries and *Escherichia coli*, 23,192 entries) with the enzyme trypsin and a maximum number of missed cleavage sites of three. Mascot was searched with a product ion mass tolerance of 0.60 Da and a precursor ion tolerance of 5.0 ppm. The carbamidomethyl of cysteine was specified in Mascot as a fixed modification. Acetylation of the protein N-terminus, oxidation of methionine, pyroglutamation of glutamic acid, and phosphorylation of tyrosine, serine, and threonine are specified in Mascot as variable modifications. Scaffold (version Scaffold_5.1, Proteome Software Inc.) was used to validate MS/MS-based peptide and protein identification. Peptide identifications were accepted when established at greater than 95.0% probability. Protein identifications were accepted when established at greater than 99.0% probability and contained at least three identified peptides.

### 4.3 Negative-stain EM

We initially imaged the sample in a negative stain to assess sample purity and quality for high-resolution cryo-EM. Five microlitres of 1:1,000 of the 33 mg/ml GroEL-UGT1A sample was absorbed to the carbon substrate of mesh 600 copper grids, glow discharged for 1.5 min, and stained in the presence of 2% (w/v) ammonium molybdate. Grids were then imaged with a Hitachi H-7650 Transmission Electron Microscope (Hitachi High-Technologies, Tokyo, Japan) operating with an accelerated voltage of 80 kV and equipped with a 1 × 1 K Tietz FastScan-F114 CCD camera (TVIPS, Gauting, Germany). Images were taken at 80,000× magnification, corresponding to a pixel size of 3.78 Å/pixel, with defocus ranging between 1.0– 3.0 μm.

### 4.4 Cryo-EM imaging and data collection

A fresh sample containing GroEL-UGT1A [33 mg/ml] at 1× dilution was immediately imaged after expression and purification with no freeze-thaw cycles. Three microlitres of GroEL-UGT1A in buffer (150 mM NaCl, 20 mM Tris-HCl, pH 7.5) was applied to glow discharged (1.5 min on “soft” mode) Holey carbon Quanitifoil® copper 300 mesh size R1.2/1.3 grids (Quanitifoil® Großlöbichau, Germany), and used within 30 min of discharging. The sample was then placed in a Vitrobot™ Mark IV (Thermo Fisher Scientific Waltham, MA, USA), and conditions were set to 4 °C with 100% humidity. The sample was blotted with a blot force = 1 for 3 s before the sample grid was immediately plunge-frozen in liquid ethane. The grid was then placed in a Titan KRIOS (Thermo Fisher Scientific) operating at 300 kV and equipped with a Cs corrector, and 2,951 movies within a defocus range between –1.0 and –2.0 μm were collected using the Falcon III (FEI, Netherlands) direct electron detector (DED) in counting mode at a magnification of 75,000×, corresponding to a pixel size of 0.87 Å/pixel at the specimen level (see **Supplementary Fig. 2** for imaging details).

### 4.5 Cryo-EM image processing and 2.8 Å consensus map refinement

All data were processed in RELION 3.1 [9] on a workstation containing four GeForce RTX 2080 GPUs (Nvidia Corporation, Santa Clara, California, United States) running on CUDA toolkit Version 11.0. The workstation was also equipped with 64-bit CPUs (Intel® Xeon® W-2265 CPU @ 3.50GHz) working on Linux distribution CentOS Linux 7. Motion correction was performed using RELIONs own implementation of the MotionCor2 [7] algorithm. CTF estimations of the aligned and summed micrographs were performed with CFTFIND-4.1 [8] and visually inspected for potential contamination. Laplacian of Gaussian (LoG) reference-free auto-picking resulted in 1,289,236 particles extracted with a box size of 300 pixels and Fourier cropped (referred to as ‘binning’ or down-sampled) to 3× the original pixel size, corresponding to 2.61 Å/pixel. The initial reference-free 2D classification was performed with a mask diameter of 240 Å resulting in 1,202,095 particles. These particles were examined for particle duplication (min. distance 75 Å) and subjected to 3D auto-refinement (C1), resulting in an initial reconstruction of 5.3 Å. Particles were re-extracted at the original pixel size of 0.87 Å/pixel and subjected to another round of 3D auto-refinement to give a 2.93 Å resolution map. Particles were then estimated for beam tilt, trefoil, 4^th^ order aberrations, anisotropic magnification, per-particle defocus ranges, per-micrograph astigmatism, and Bayesian polished until no improvements in resolution and B-factor were reported. The final consensus reconstruction (C1) is reported at 2.8 Å resolution by an FSC cut-off of 0.143 with an improved sharpening B-factor of –91 (originally –121.9). See **Supplementary Fig. 2** for the refinement workflow and details and **Supplementary Fig. 3** for FSCs and orientation distribution.

### 4.6 Polyshape masking and the separation of SER/DOR complexes from the consensus map

The polyshape soft mask was generated using a combination of tools from the SPIDER suite [41] and UCSF Chimera-X [42]. We generated two density volumes to be added together. A cylindrical volume was generated by the Create Model Volume tools in SPIDER with a height of 75 pixels and a radius of 25 pixels. The second volume was created in Chimera-X using the “molmap” command from the fitted atomic coordinates of three GroEL subunits (two contiguous and one transversely across the ring) that only contained residues 197–279 and 298– 319. These two volumes were added together using the “volume add” command in Chimera-X and imported into the RELION pipeline with the Mask Create operation (extend: 3 pixels; soften: 3 pixels) (**Supplementary Fig. 12**).

Two rounds of masked 3D classification were performed with the polyshape mask positioned over the ring cavity of the 2.8 Å consensus map (**Fig. 2**). The mask was placed on the bottom and top cavity in turn, classified without alignment, i.e., without performing an orientational search, and particles were selected based on the presence or absence of UGT1A density within a ring cavity (referred to as focus-cleaning) (**Supplementary Fig. 13**).

### 4.7 Subunit-focused classifications and visualisation in the context of GroEL

Generating a soft mask around the GroEL protomer (subunit) from both UGT1A-empty and UGT1A-occupied ring complexes, SER(2)-GroEL-UGT1A and DOR-GroEL-UGT1A, respectively, was performed in Chimera-X using the molmap command (resolution = 7 Å) from fitted atomic coordinates. The resulting density volumes were Gaussian filtered (σ = 3) and imported into the RELION pipeline with the Mask Create operation (extend: 3 pixels; soften: 1 pixel).

The resulting protomer mask was applied directly to the DOR-GroEL-UGT1A complex (C1) and subjected to a single round of focus classification (*K* = 10). Because of the relatively low number of particles in the SER(2)-GroEL-UGT1A (C1) map, we symmetry expanded the particle stack using the relion_particle_symmetry_expand command with D7 symmetry from a unit cell over the UGT1A empty ring cavity only (**Supplementary Figs. 5** and **6**). From this D7 symmetry expanded particle stack, a soft mask around a single subunit was applied (as described above) and subjected to a single round of focus classification (*K* = 10).

### 4.8 Flexible ATP Docking

Flexible docking was performed using Glide software Maestro (Schrodinger Inc). The ER and OR subunits were pre-processed, and the H-bond network was optimized and minimized as receptors using the protein preparation wizard with default settings. There were no reported stereo clashes of overlapping atoms.

Preparation of ATP (taken from PDB ID: 3WVL) was performed by the Ligand Preparation (LigPrep) wizard. The ligand geometries were optimized and correct chiralities were determined using the OPLS _2005 forcefield (pH 7.0 +/– 2.0 from the input 3D structure). Two ATP molecules were generated, the original 3WVL-ATP and a chiral alternative conformation based on the 3D architecture. Both ATP molecules were then prepared for ligand docking to GroEL. Next, a 30 Å^3^ grid box was generated from a centroid of specified residues (87, 151, 495, and 33) that best define the presumed ATP binding pocket of GroEL.

Lastly, ligand docking was performed using Glide. The two ATP molecules were used for docking into the pre-processed 30 Å^3^ grid, and box and docking precision was set to Standard Precision (SP) with flexible ligand sampling. The number of poses per ligand was set to 50 to generate a variety of docking poses. The resulting docking energy scores were recorded and filtered to RMSDs of < 4.5 Å from the reference 3WVL ATP positioning (see the legend to **Supplementary Fig. 11** for details).

### 4.9 Modelling and refinement

All models were initially rigid fit to the EM coulomb density in Chimera-X v1.2.5 [43], and atomic coordinates were modelled in Coot v0.9.5 [44], with final refinements and validation performed in Phenix v1.19.2-4158 [45, 46] using the phenix.real_space_refinment tool. Initially, the X-ray crystallographic model PDB ID: 2NWC was rigidly fit into our 2.8 Å consensus map. Side-chain residues were manually modelled in Coot and passed iteratively through Phenix refinement and Coot reducing Ramachandran outlier, rotamer, and geometry minimisation. The resulting atomic coordinates were used in all subsequent structure refinements. See **Supplementary Tables 1–5**.

## Data availability

The 3D cryoEM density maps are deposited in the Electron Microscopy Data Bank (EMDB) with the following accession numbers, 2.8 Å consensus map: EMD-33349, 2.7 Å DOR-GroEL-UGT1A: EMD-33350, 3.2 Å SER(2)-GroEL-UGT1A: EMD-33351, 3.2 Å OR4-GroEL-UGT1A (with clearest UGT1A density in cavity): EMD-33352, and subunit conformations ER1: EMD-33353, ER2: EMD-33354, OR1: EMD-33355, OR2: EMD-33356, OR3: EMD-33357, and OR4: EMD-33358.

The corresponding atomic coordinates are deposited in the Protein Data Bank (PDB) with the following IDs, 2.8 Å consensus map: 7XOJ, 2.7 Å DOR-GroEL-UGT1A: 7XOK, 3.2 Å SER(2)-GroEL-UGT1A: 7XOL, 3.2 Å OR4-GroEL-UGT1A (with poly-alanine peptide): 7XOM, and subunit conformations ER1: 7XON, ER2: 7XOO, OR1: 7XOP, OR2: 7XOQ, OR3: 7XOR, and OR4: 7XOS. Raw movies of all datasets have been deposited in the Electron Microscopy Public Image Archive (https://www.ebi.ac.uk/pdbe/emdb/empiar/) with accession codes **XXXXX**.

## Author Contributions

E.M. conceived the project; K.S., T.M., J.T. and E.M. designed the experiments; T.M., T.H. and E.M. expressed and purified proteins; T.Kawamura performed LC-MS/MS analysis; K.S. carried out cryo-EM experiments with the support of N.M., T.Kato, K.I. and T.N.; K.S. and E.M. prepared atomic models; K.S. and D.M.S. conducted simulation analysis; K.S., J.T. and E.M wrote the manuscript, with input from all authors.

## Acknowledgements

We thank Zuben P. Brown and Eiichiro Ono for helpful discussion, Mika Hirose and Junichi Kishikawa for assistance of cryo-EM data collection, Takanori Hongo, Taichi Hayashi, Taisuke Nakayama and Reiko Sato for assistance of protein production, and Yoshihiro Maruo for provision of the UGT1A gene. We acknowledge funding from the Platform Project for Supporting Drug Discovery and Life Science Research (BINDS) from the Japan Agency for Medical Research and Development (AMED) (21am0101075 to J.T., 22ama121025j0001 to D.M.S. and JP21am0101066 to G.K.); the Japan Society for the Promotion of Science (JSPS), KAKENHI (JP19K06513 to T.M.); and to E.M., the Asahi Glass Foundation; the Novartis

Foundation (Japan) for the Promotion of Science; a Grant-in-Aid for Scientific Research on Innovative Areas from the Ministry of Education, Culture, Sports, Science, and Technology (MEXT) (19H05780); the Japan Science and Technology Agency (JST), PRESTO (JPMJPR17GB); and a project, JPNP18016, commissioned by the New Energy and Industrial Technology Development Organization (NEDO).

