## Supplementary Fig for "Three-dimensional motions of GroEL during substrate protein recognition"

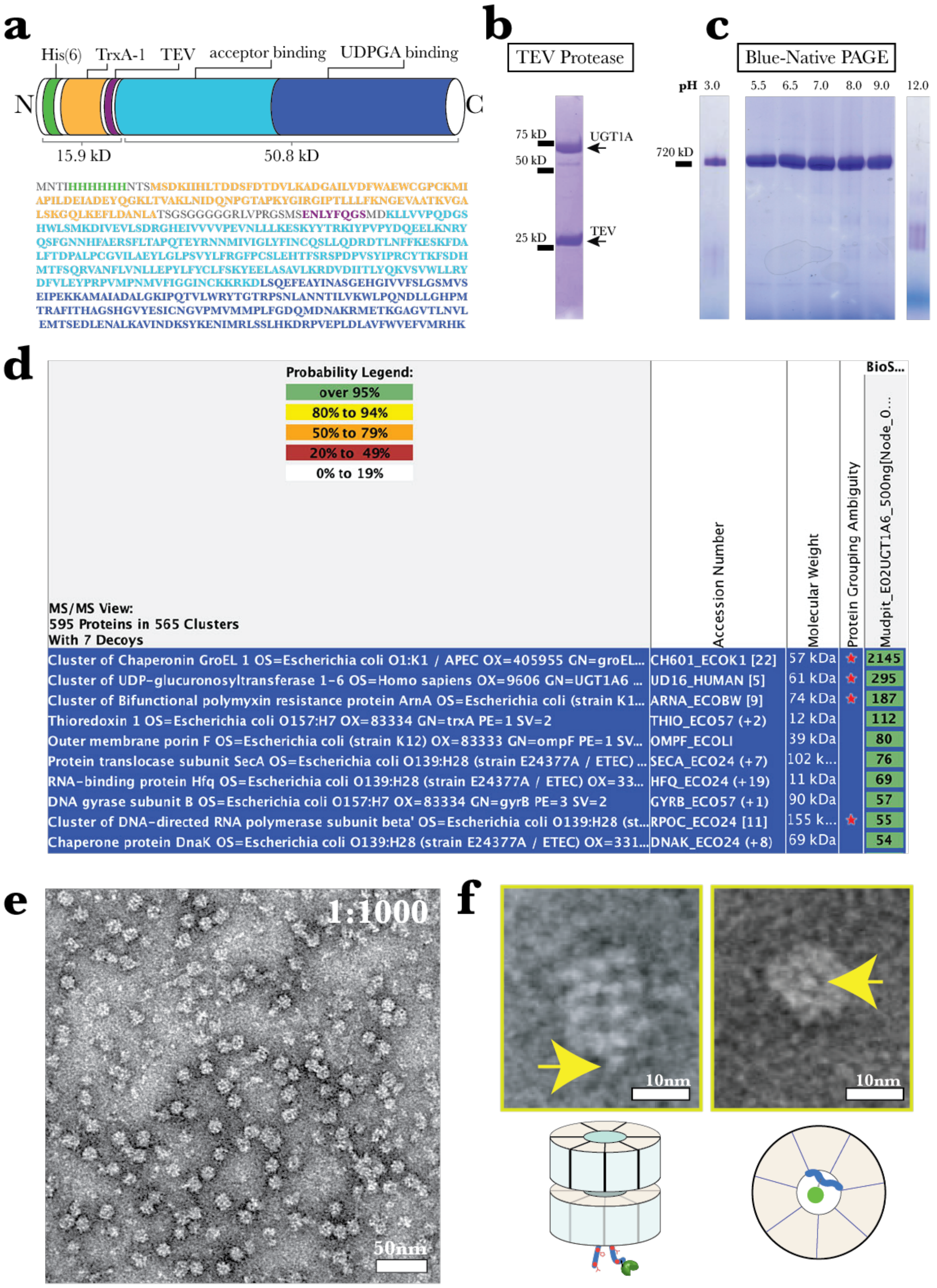

**Supplementary Fig. 1. Expression, identification, and visualisation of GroEL-UGT1A.**

**a,** Schematic of the UDP glucuronosyltransferase 1A (UGT1A) expression sequence and construct with His$_6$-tag (green), thioredoxin (TrxA) tag (yellow), TEV cleavage site (purple), and UGT1A acceptor binding and UDPGA binding domains (light and dark blue, respectively). **b**, Separation of TEV protease and purified

UGT1A by SDS-PAGE. Incubation with TEV protease (~26 kDa) did not change the mobility of the UGT1A band (~70 kDa), indicating that cleavage did not occur. **c**, Blue native PAGE of the elution from nickel affinity chromatography shows a clear band at ~720 kDa over pH values pH 3.0–9.0, with no band observed at pH 12. **d**, LC-MS/MS proteomic analysis performed on the purified UGT1A sample identified large amounts of a 57 kDa *Escherichia coli* chaperonin GroEL within the purified sample. **e**, Negative-stain evaluation of a sample containing 33 mg/ml GroEL-UGT1A diluted 1:1,000 (scale bar: 50 nm). GroEL-UGT1A particulates are distributed evenly in the field of view and fixed in multiple orientations with no visible aggregation. **f**, Side and top views of a GroEL-UGT1A complex (scale bar: 10 nm). The yellow arrow points to an extra density corresponding to UGT1A visibly bound to a GroEL ring.

**TITAN Imaging Conditions**

| | |
|---|---|
| Microscope | Titan Krios G3 |
| Voltage (kV) | 300 |
| Magnification | x75,000 |
| Probe | Nanoprobe |
| Detector | Falcon III |
| Mode | Counted |
| Data collection software | EPU |
| Electron exposure (e–/Å2) | 40 |
| Exposure time (seconds) | 36.9 |
| Number of fractions (Nr) | 50 |
| e-/px/s | 0.89 |
| e–/Å2/s | 1.08 |
| Spot size | 9 |
| Intensity | |
| C2 apperature | 50 |
| Defocus range (µm) | -1.2 ~ -2.0 µm |
| Image size (px) | 4096 x 4096 |
| Spherical Aberration (mm) | hardware corrected |
| Pixel size (Å) | 0.87 |

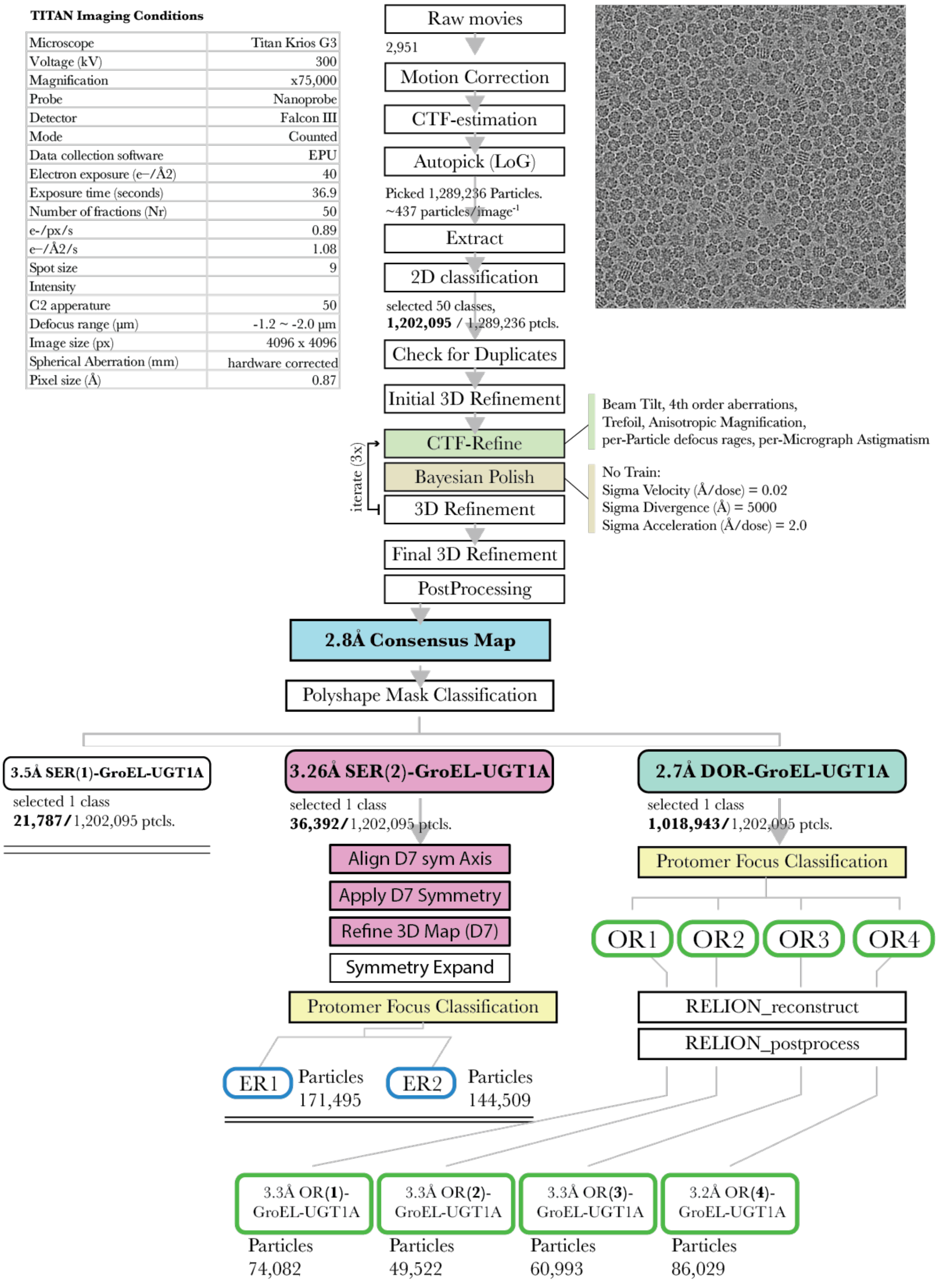

### Supplementary Fig. 2. Imaging and refinement processing pipeline.

Titan imaging conditions and the single-particle processing pipeline for all reconstructions.

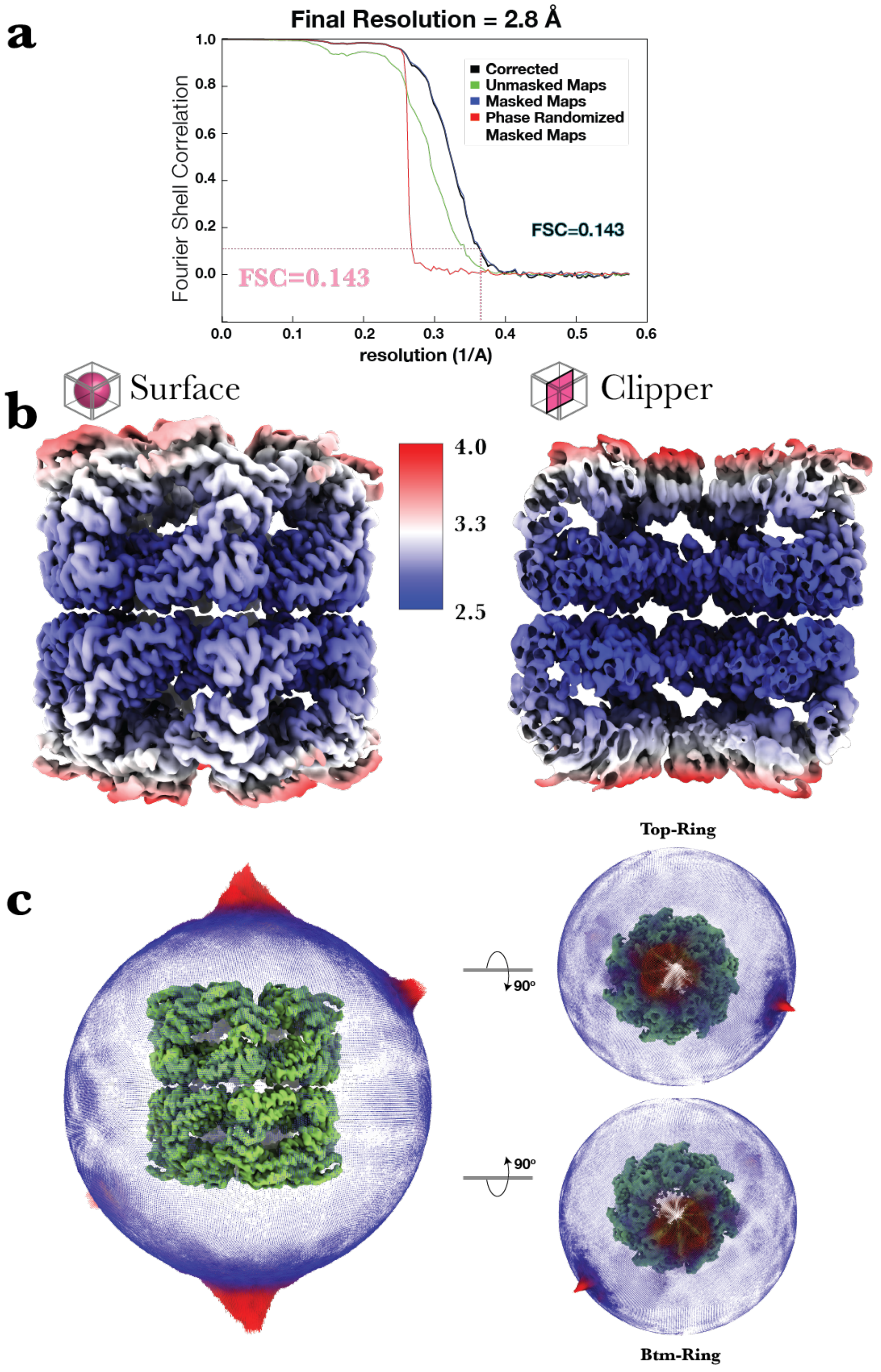

**Supplementary Fig. 3. FSC, local resolution and particle distribution.**

**a**, The consensus map's FSC curves. Global resolution is 2.8 Å (FSC cut-off = 0.143). There is no evidence of overfitting (red curve) or signal enhancement caused by the convolution effect (black curve), and the correlation between masked (blue curve) and unmasked (green curve) half maps is good. **b**, The consensus reconstruction shown in surface and clipper views is coloured by local resolution: blue-white-red representing 2.6–3.0–3.5 Å. **c**, Euler angular distribution of the refined consensus map particles. Each dot serves as a visual cue to show from which point of view or Euler angle a particle was assigned during refinement. The number of particles assigned to that angle is indicated by the colour (blue to red) and length. From that Euler angle, red and long have many particles of that view, whereas blue and short have fewer. Thus, visual inspection of the particles making up a reconstruction was assessed. This illustration shows a uniform distribution of assigned particle Euler angles, making a complete and nearly unbroken sphere of views with many particle images representing top, bottom, and orthogonal views (referred to as 'hot spots' or orientation bias).

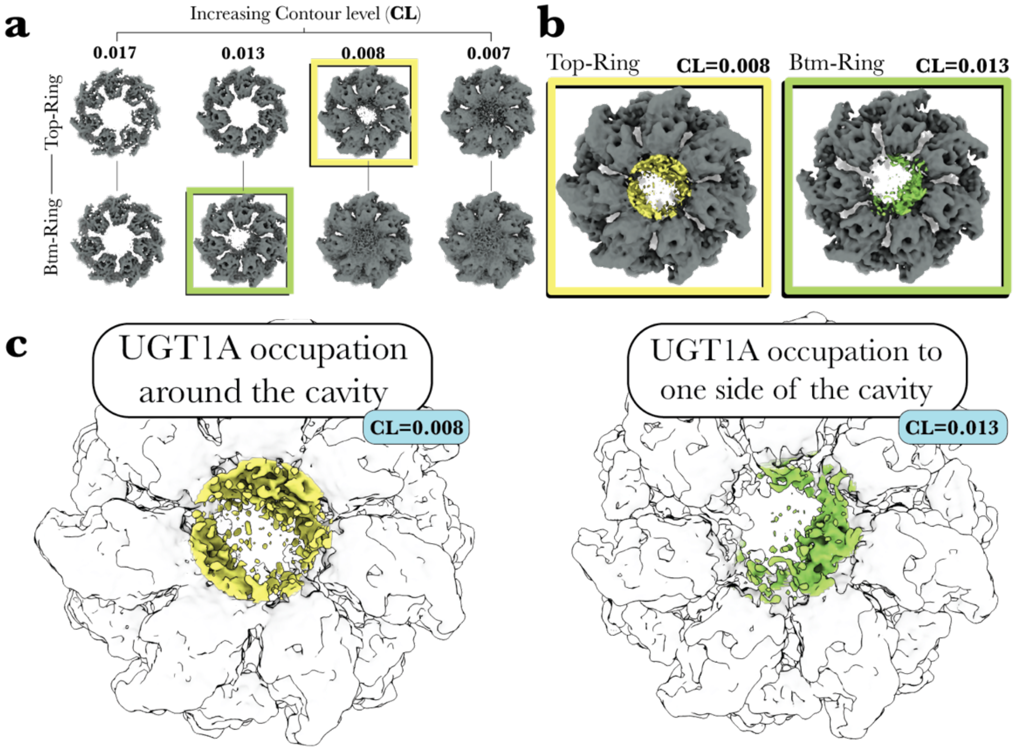

**Supplementary Fig. 4. Different UGT1A occupation motifs within the consensus map rings.**

**a**, Within the top and bottom rings, changing the contour level (CL) of the consensus map density volume yields distinct substrate density occupation motifs. The flexible UGT1A in the top-ring (yellow square) emerges at lower contour levels than in the bottom-ring (green square). **b**, UGT1A occupation between the rings was compared by selecting top and bottom ring images from different contour levels. The UGT1A density is coloured yellow and green and occupies the top and bottom rings (dark grey). **c**, GroEL (white silhouette) and UGT1A densities (coloured). Different occupation patterns are produced by adjusting the contour level of the two volumes until UGT1A is visible in the ring. The bottom ring is asymmetrically occupied by UGT1A, whereas the density of the top ring is dispersed around the cavity. This result can only occur if, during refinement, particles were aligned with respect to the UGT1A occupation pattern in the bottom ring regardless of substrate occupation in the top ring, and it is the first observation that asymmetric binding in one ring has no influence on substrate binding in the other ring.

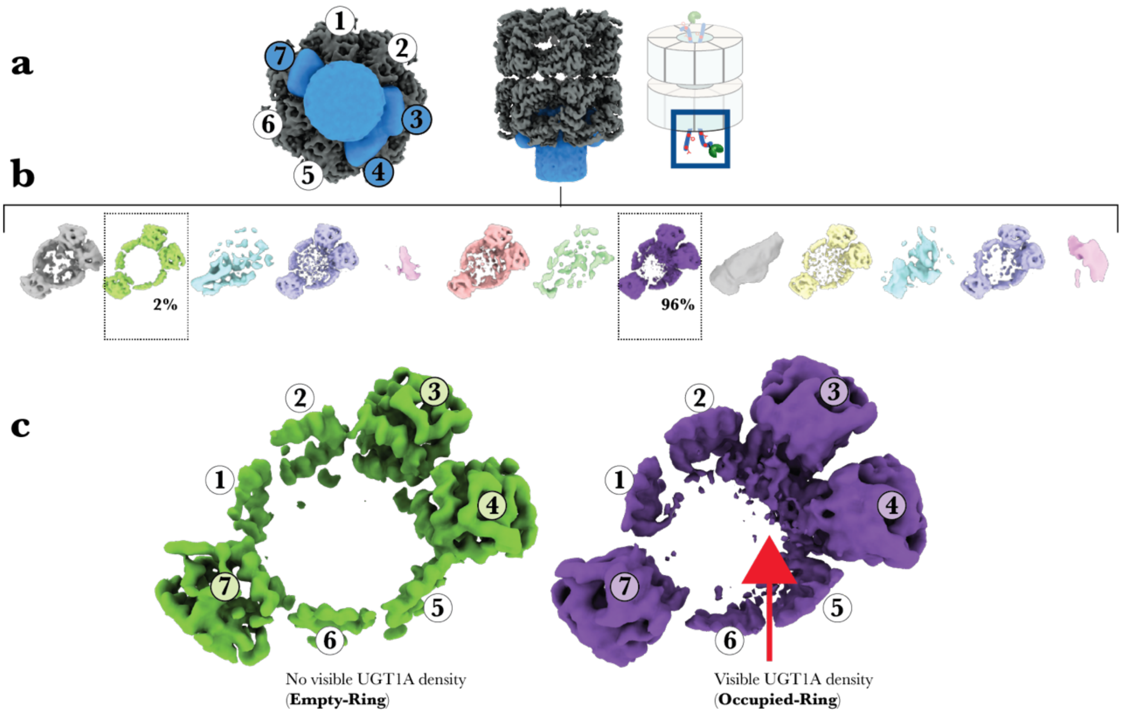

**Supplementary Fig. 5. UGT1A Asymmetric binding within a GroEL ring and identifying the substrate-empty class of particles.**

**a**, Polyshape mask (blue) is positioned on the bottom ring of the consensus map (grey), and subunits of that ring are numbered 1–7. Apical domains of three subunits (3, 4, and 7), including the cavity region, are covered by the polyshape mask for separation by focused mask classification. **b**, Particle images are separated into two discrete 3D volumes (green and purple). A minority class volume (2%) and a dominant class volume (96%) after one cycle of focus classification into fourteen classes. **c**, Two selected volumes in close-up with apical domain positions labelled 1–7. The UGT1A density (red arrow) asymmetrically forms to one side of the ring cavity, contacting 2–3 apical domains in the dominant volume. However, in the ring cavity of the minority volume, there is no UGT1A-occupying density, showing that this classification method can distinguish between occupied and empty states of the GroEL ring.

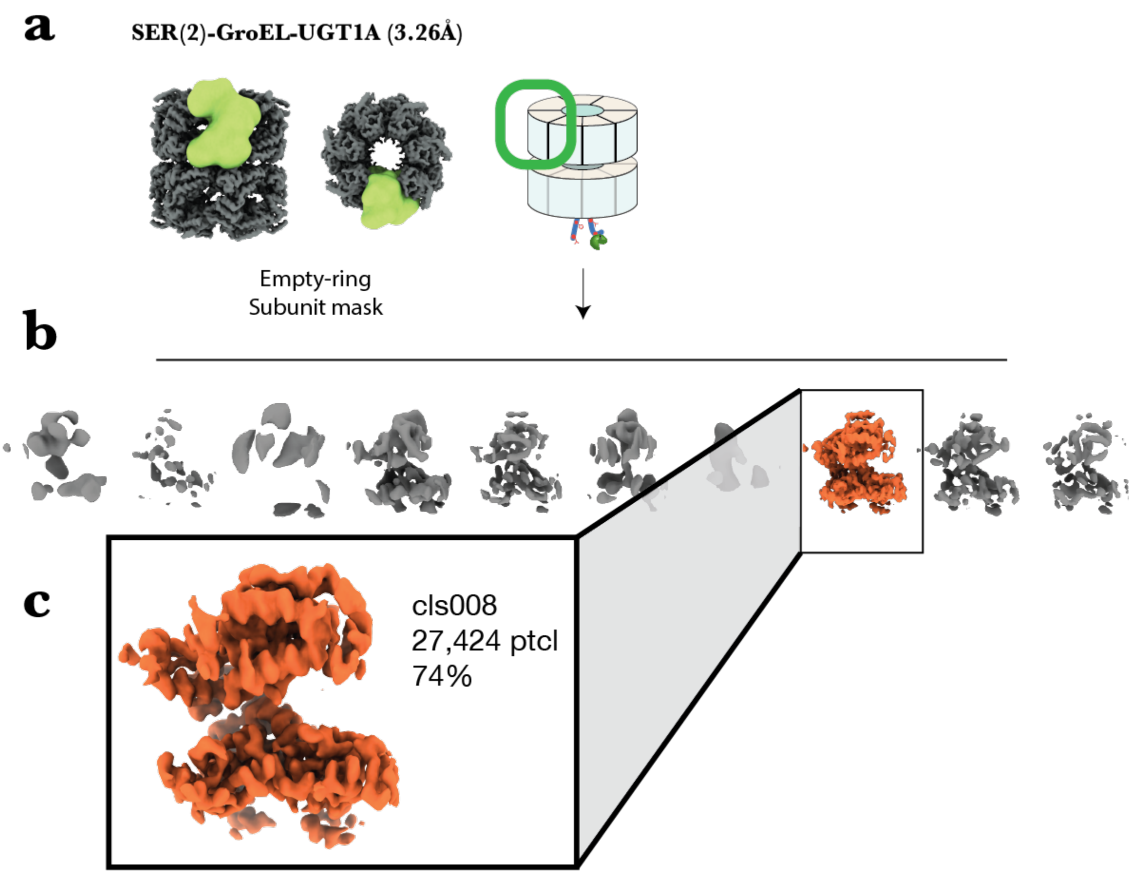

**Supplementary Fig. 6. Initial focus classification of the SER(2)-GroEL-UGT1A asymmetric reconstruction (C1).**

**a**, A soft mask (green) is applied to a single subunit in the bottom ring of SER(2)-GroEL-UGT1A (grey). **b,c**, One discrete class (orange) was classified from a single round of focus classification ($K$ = 10), representing 74% of the total image population.

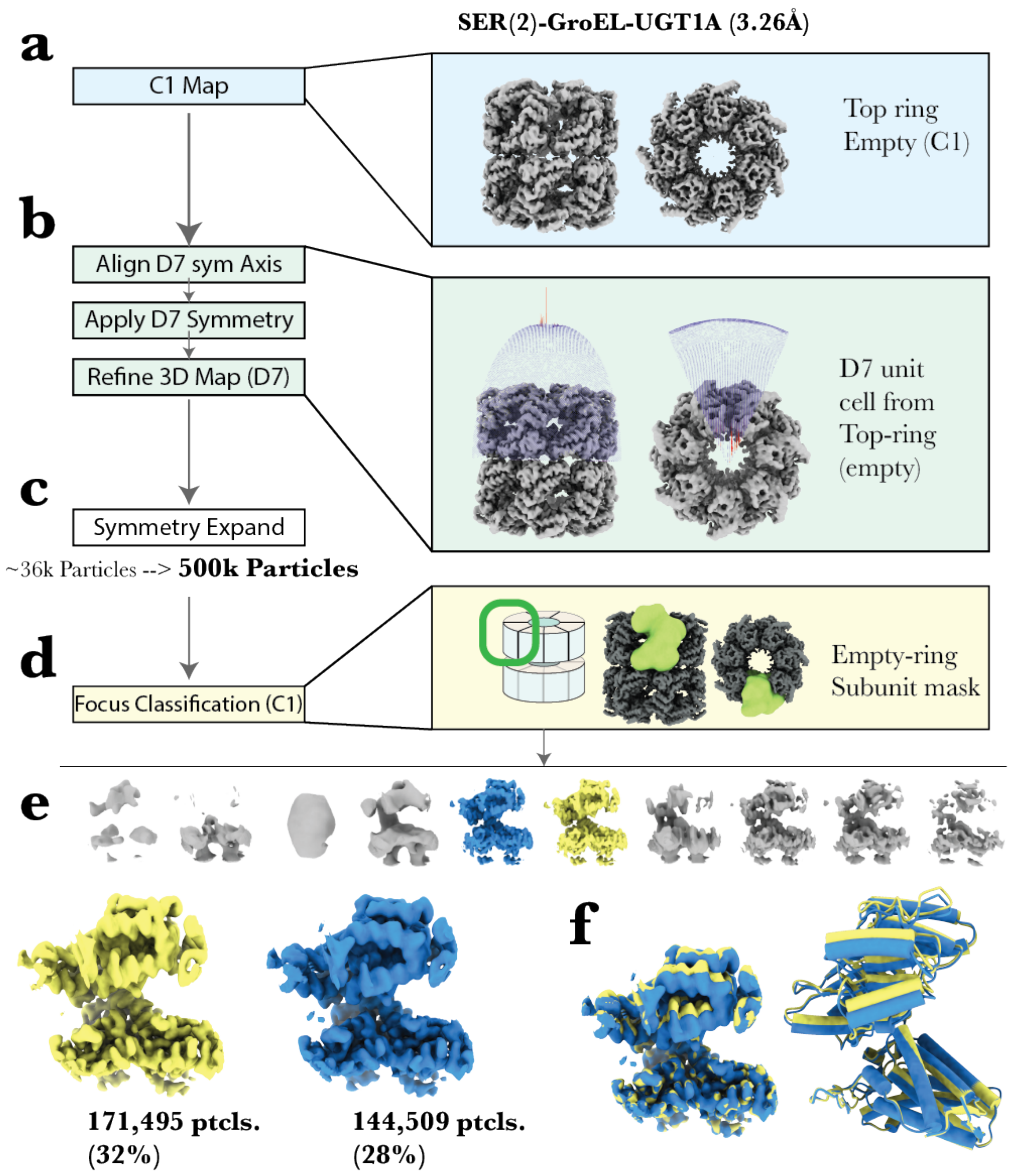

**Supplementary Fig. 7. Symmetry expansion of the SER(2)-GroEL-UGT1A density map for focus classification of an empty ring protomer.**

**a**, The refined C1 (no symmetry) SER(2)-GroEL-UGT1A density map (blue box) is shown as side-on and top views. The top ring of the complex is absent of any UGT1A density. **b**, Aligning the volume density along the D7 symmetrical axis positions the top ring (empty) as the ring used for unit cell point group symmetry. After aligning and applying symmetry (D7) to the density volume and all particle images, the map and particle images were refined by imposing D7 symmetry. The unit cell used in this point group symmetry can be seen here as a 'wedge' of blue dots (particle distribution plot). **c**, Symmetry expansion of a D7 imposed refined particle stack increases the number of particle projections as a multiple of the number of symmetry elements in a point group symmetry used. In this case, 14×, therefore, 36k (C1) particle projections increase to ~500k (D7) particle projections. **d**, A soft mask is placed over a single subunit (green), and one round of focus classification (**e**) yields two discrete 3D density volumes (yellow and blue), representing 32% and 28% of the image population, respectively. These conformations were designated ER1 and ER2. **f**, Superimposition of ER1 and ER2 density volumes and their refined atomic coordinates (cylinders and stubs).

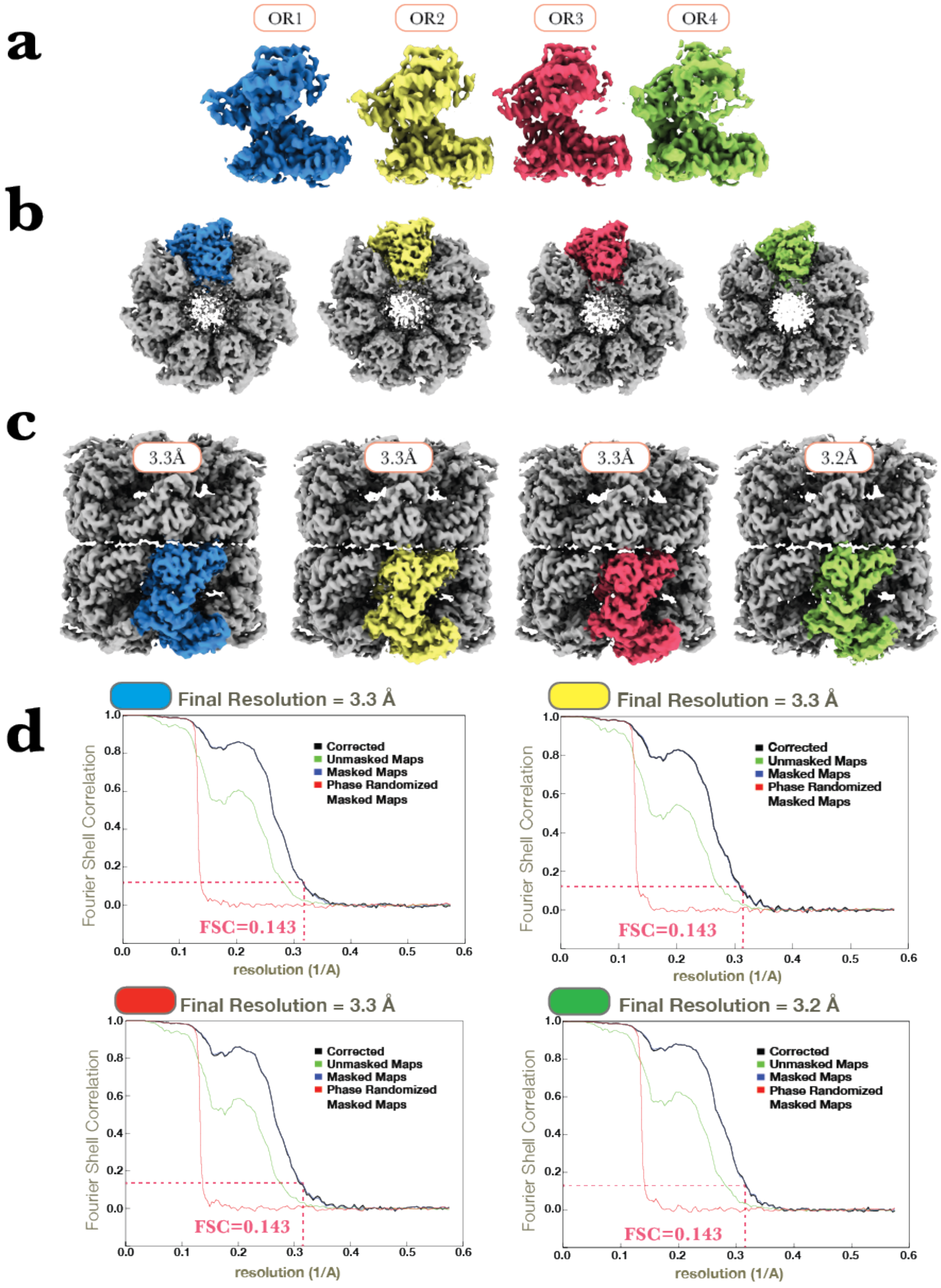

**Supplementary Fig. 8. OR(1–4)-GroEL-UGT1A.**

**a**, The focus maps (OR1–4) are coloured blue, yellow, red, and green, respectively. **b**, The focus map 3D volumes are coloured according to the conformer discrete maps in (**a**) and are observed in the bottom ring of GroEL-UGT1A when viewed face-on. In all three reconstructions, the UGT1A density is observed asymmetrically occupying a GroEL ring and contacting 2–3 subunits. **c**, The focus maps were reconstructed and refined without the soft mask used in focus classification and are shown here in the context of the GroEL binary ring complex. **d**, All maps were reconstructed according to the gold-standard reconstruction method, and the global resolution was calculated using an FSC cut-off of 0.143 (dashed pink line). The masked and unmasked half maps' FSC curves (blue and green curves) are both healthy, with no symptoms of overfitting (red curve) or enhanced signal caused by the convolution effect (black curve).

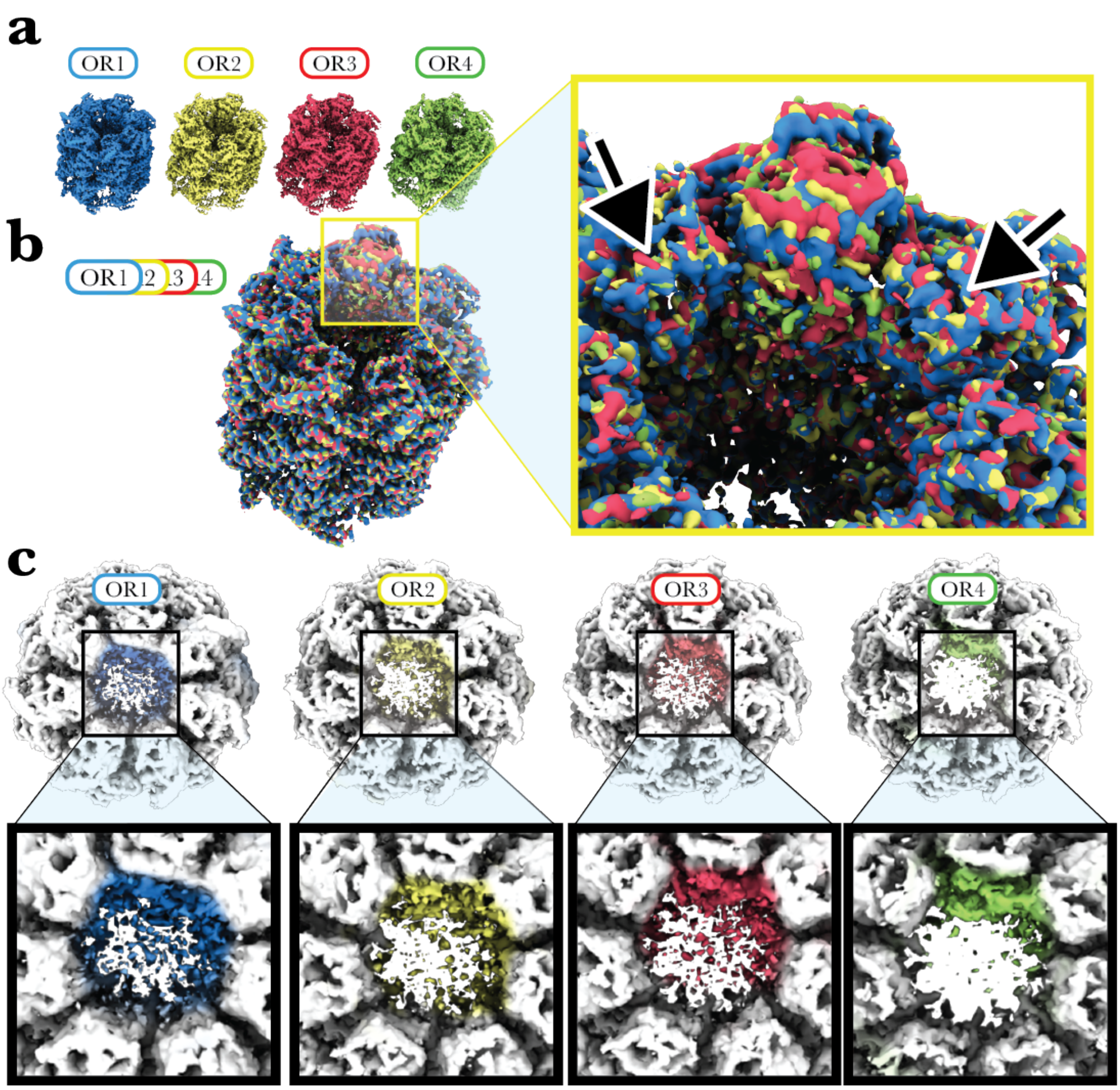

**Supplementary Fig. 9. Asymmetric ring occupation motif and unaffected subunit neighbours.**

**a**, The final refined OR(1–4)-GroEL-UGT1A complexes coloured by focus classification of the 3D volume class origin (Fig. 3). **b**, 3D volumes of OR(1–4)-GroEL-UGT1A complexes superimposed. Except for the conformation of the OR protomer derived from focus classification (magnified yellow box), which can be seen breaking the ring symmetry because of elevation in the apical domain, the overall fit of the four maps shows minimal-to-no deviations in the main-chain and side-chain corresponding densities. Neighbouring protomers on either side of this protomer show no evidence of apical domain elevation (black arrows). **c**, The asymmetric occupation of the UGT1A density (coloured) within a GroEL ring is shown in the four OR(1–4)-GroEL 3D volumes (white). Density formation in UGT1A contacts 2–3 subunits localised around a core protomer, and density formation becomes more prominent as the elevation in the apical domain increases.

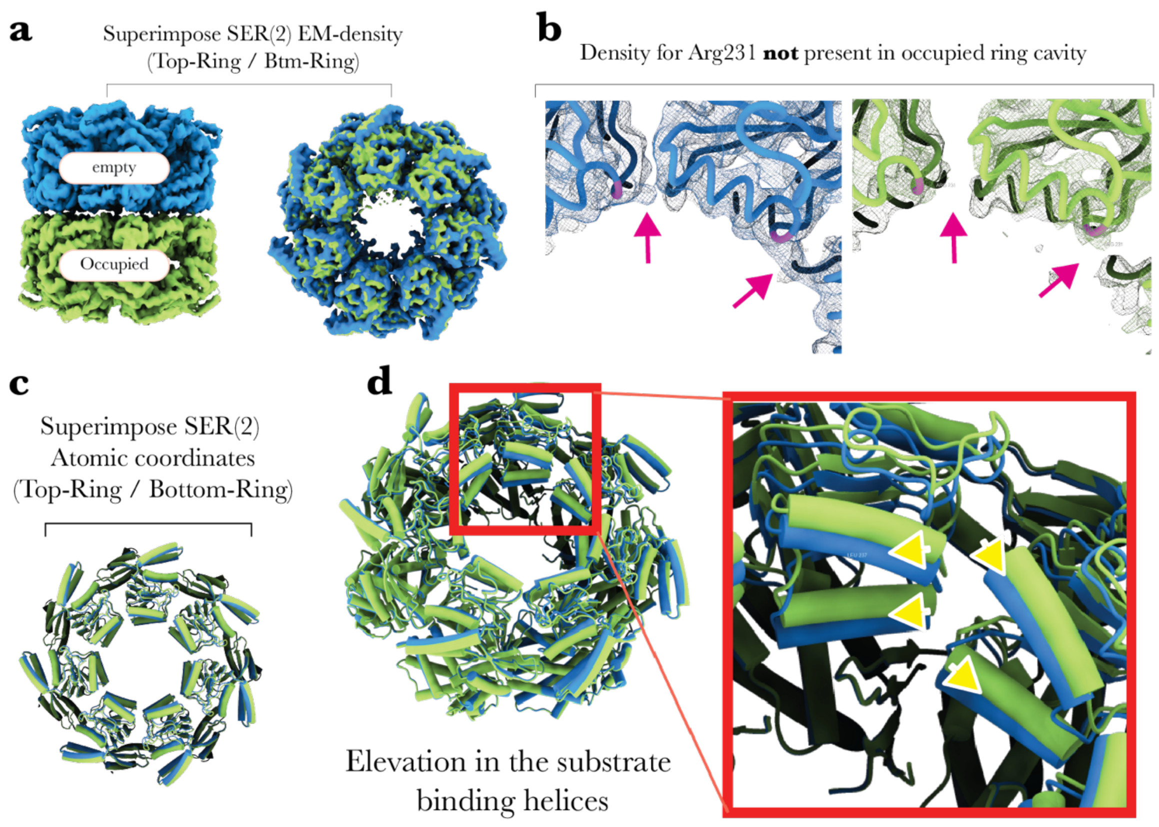

**Supplementary Fig. 10. Loss of Arg231 density and substrate binding site elevation.**

**a**, The top and bottom rings from the SER(2)-GroEL-UGT1A density volume are superimposed. The GroEL ring is either occupied with UGT1A density (green) or empty (blue). From a top-view vantage, the occupied ring appears to exhibit a slight cavity ring expansion. **b**, When the atomic coordinates (shown in liquorice) were fitted and refined to the SER(2) complex density volume (surface mesh), the density corresponding to side-chain Arg231 (pink liquorice) was evident in the empty ring but not in the occupied ring (pink arrows). **c**, Superimposition of the atomic coordinates from the top (empty) and bottom (occupied) rings are depicted as cylinders and stubs. **d**, The occupied ring (green) exhibits a uniform elevation of substrate binding helices H and I (yellow arrows) lining the GroEL ring cavity.

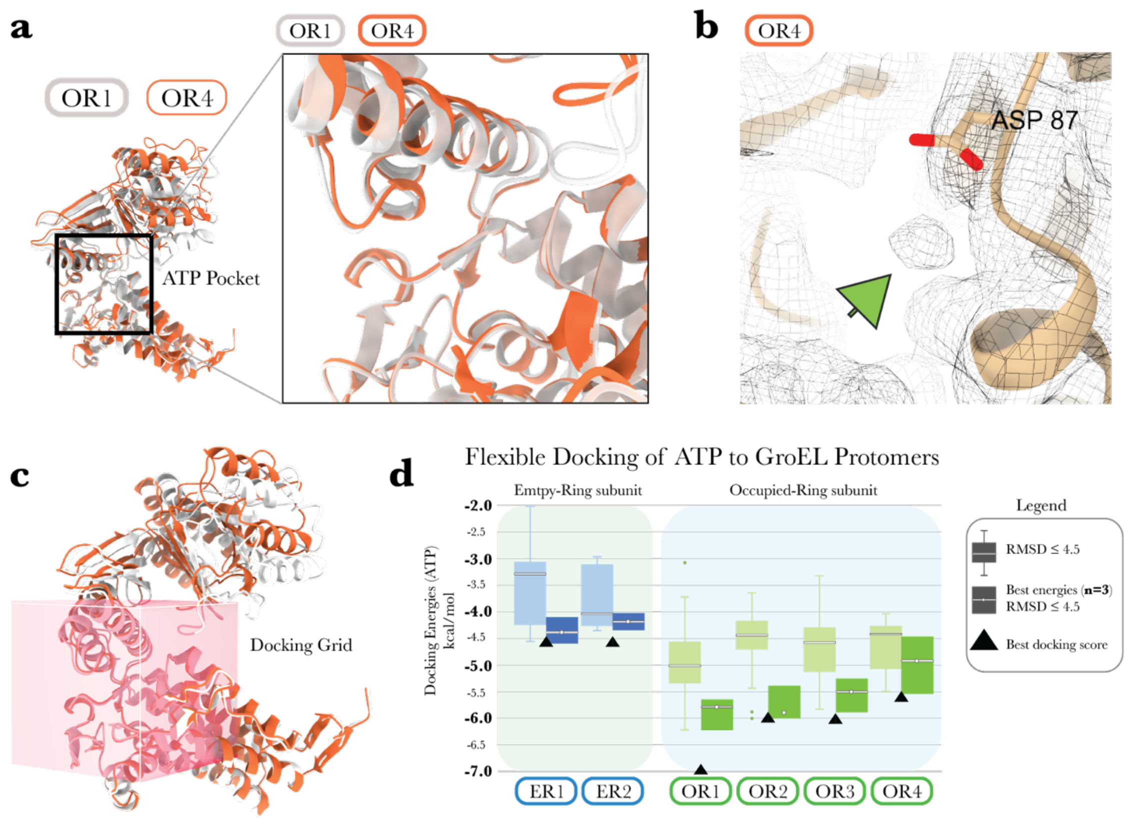

**Supplementary Fig. 11. Flexible docking simulation of ATP to empty and occupied ring protomers.**

Flexible movements in the nucleotide-binding pocket and ATP docking energies. **a**, ER1 (white) and OR4 (orange) atomic coordinates are superimposed and shown in ribbon representation. Significant structural variations exist in the ATP binding pocket that may be allosterically coupled to the substrate binding helices in the apical domain (black squares). **b**, OR4 volume density (grey mesh) and fitted atomic coordinates (shown in ribbon representation). Near one of the ATP hydrolysing residues in OR4 strong density has formed (green arrow) and is positioned conspicuously where Asp87 coordinates $Mg^{2+}$ in other published structures. This density is present only in the OR4 density volume. **c**, The 30 Å$^3$ ligand docking grid box (pink cube volume) centred on Asp87, Asp495, Ser151, and Pro33 was used for flexible docking of ATP into the empty ring protomer (ER1–2) and occupied protomer OR1–4. Shown positioned over ER1 and OR4 is the grid box volume used to calculate docking energy scores for all protomers. **d**, Docking scores (kcal/mol) of ATP to each protomer. The whisker box plot shows the docking energies filtered by RMSD values ≤ 4.5 between atom pairs of docked ATP to the GroEL protomer and ATP from PDB ID: 3WVL. The box plot displays the top docking energies from those filtered by RMSD ≤ 4.5 ($n$ = 3), and the black triangles represent the best docking energies unfiltered.

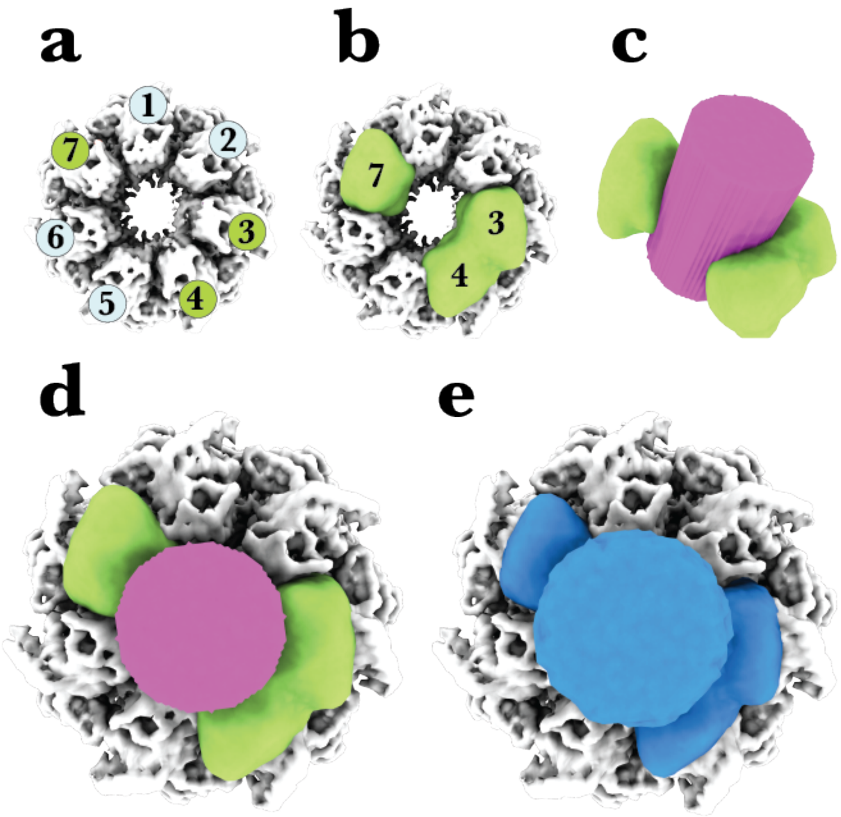

**Supplementary Fig. 12. Polyshape mask creation.**

**a**, Bottom ring of the GroEL-UGT1A density map, with subunits numbered 1–7. **b**, After rigid fitting the atomic coordinates to the GroEL-UGT1A density, residues 197–279 and 298–319 from subunits 7, 3, and 4 were transformed to a density volume using the "molmap" command corresponding to 5 Å. The resulting tri-subunit density (green) was then smoothed by filtering the pixel intensities by a σ value of 3 using the "vop gaussian" command in chimera X. **c**, In SPIDER, a cylinder volume (pink) was created and manually positioned over the centre of the cavity, overlapping with the tri-subunit density volume. **d**, Illustration of the uncombined volumes (tri-subunits and cylinder) before using the "volume add" command to merge them into a single continuous density. **e**, The cylinder and tri-subunits have now been joined to form a single, continuous soft mask, referred to as the "polyshape mask" volume. This volume is then resampled over the GroEL-UGT1A ring cavity after the map edges have been 'softened' by filtering the pixel intensities of the mask by a σ value of 3.

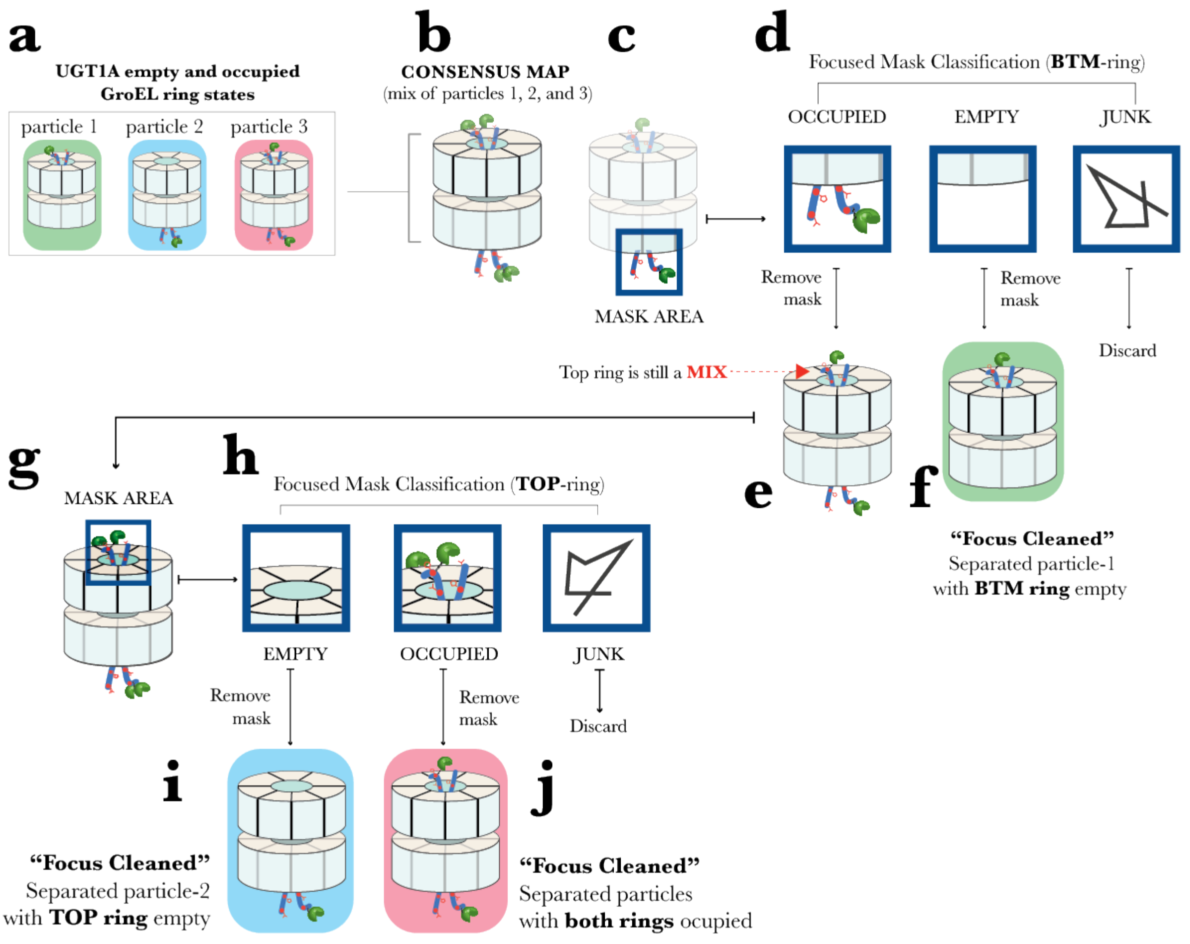

**Supplementary Fig. 13. The focus cleaning method.**

The schematic shows how to distinguish various GroEL molecules (particle images), with different substrate (UGT1A) occupations, from a consensus map with fixed particle image orientations (Euler angle assignments) that cannot be modified. **a**, Three examples of GroEL-UGT1A particle images that exhibit different substrate occupation states (compositionally heterogeneous) present in the image population. In this example, UGT1A occupies a single ring (particle-1 and -2) or occupies both rings (particle 3). **b**, A consensus map reconstruction of fixed orientation particle images from (**a**). The illustration represents a map where all three particle image examples from (**a**) have been aligned into a consensus reconstruction and are indistinguishable, giving the false impression of a GroEL-UGT1A complex with both rings occupied. **c**, To examine and separate any compositional heterogeneity in this consensus map, a soft mask is initially placed over the bottom cavity region (blue square) for particle image separation by focus mask 3D classification (without alignment). **d**, The separation of particle images by focus classification separates particles into discrete classes. In this example, there are three possible classes. An 'occupied' 3D volume class of particle images where UGT1A occupation is detected in the cavity. An 'empty' 3D volume class of particle images where no detectable UGT1A density is present, and a 'junk' class of particle images where no reasonable 3D volume can be visualised and is therefore excluded from further processing. **e**, A reconstruction of the GroEL-UGT1A density volume without the mask from occupied class particle images. The resulting GroEL-UGT1A density volume in the bottom ring has been 'cleaned' of compositional heterogeneity, i.e., any particle images with an empty bottom-ring or junk.

**f**, A reconstruction of the GroEL-UGT1A density volume without the mask from empty class particle images. Because purification of the GroEL-UGT1A complex was performed by nickel affinity of the $His_6$-tag in the UGT1A complex (see Section 6.2), there must be at least one ring with UGT1A occupation. Therefore, this reconstruction must represent particle-1 from (**a**). **g**, The 3D volume of GroEL-UGT1A reconstructed from the occupied class of particle images is selected for the second round of separation of heterogeneity by focus mask classification. However, the mask is now positioned over the top ring of this volume to identify and separate empty, occupied, and junk classes of particle images from the top-ring (**h**). **i**, A reconstruction of the GroEL-UGT1A density volume without the mask from empty class particle images. Both rings of this reconstruction have been expunged of any compositional heterogeneity by focus mask classification (focus-cleaned). By selecting bottom-ring occupied particle images in the first round of classification and empty top-ring particle images in the second round of classification, the resulting volume accurately reflects particle-2 in (**a**). **j**, A reconstruction of the GroEL-UGT1A density volume without the mask from occupied class particle images. By selecting bottom-ring occupied particle images in the first round of classification and occupied top-ring particle images in the second round of classification, the resulting volume accurately represents particle-3 in (**a**). Using this logical approach, the separation of compositional heterogeneity by focus mask classification is performed without altering the assigned Euler angles (particle image orientation) fixed during consensus map reconstruction.

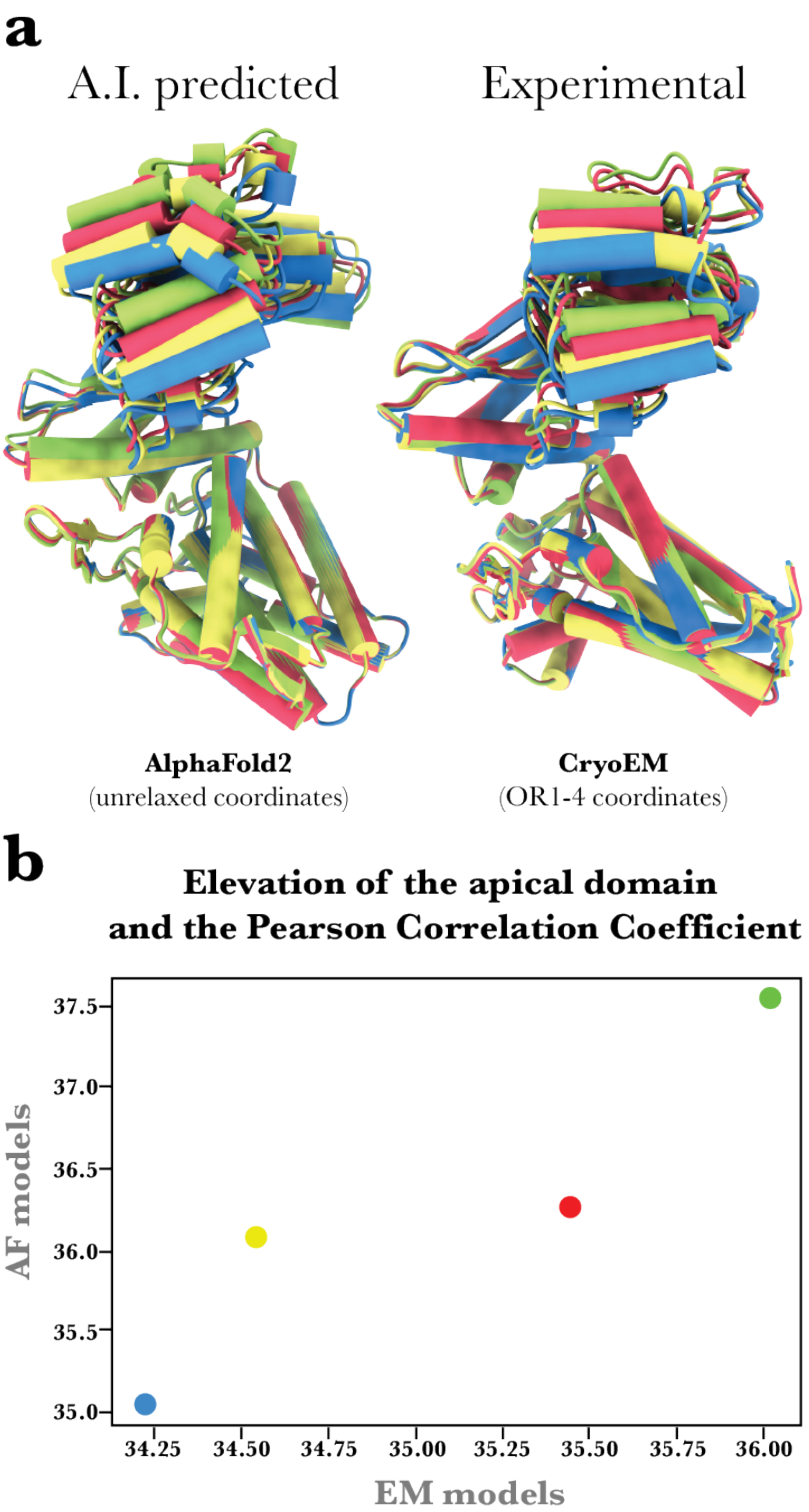

**Supplementary Fig. 14. Predicted structural dynamics of a GroEL subunit.**

AlphaFold2 (AF2) subunit dynamics prediction of the large apical domain elevation correlates with experimentally derived cryo-EM atomic coordinates. **a**, The AF2 predicted coordinates next to the EM derived coordinates from focused mask classification. **b**, Comparison between apical domain elevation in AF2 predicted models (*y*-axis) and cryo-EM atomic models (*x*-axis) using the Pearson Correlation coefficient. The Pearson Correlation coefficient is 0.925. In contrast, when the order of the AF models was shuffled randomly, the correlation dropped to 0.040 +/– 0.586.

**Supplementary Table 1. 2.8 Å consensus map data processing and refinement.**

| **Data processing** | **2.8Å Consensus Map** |
|---|---|
| Number of micrographs | 2951 |
| Number of frames per-micrograph | 50 |
| Applied dose per frame (e–/Å2) | 0.798 |
| Final particle images | 1,202,095 |
| Symmetry imposed | C1 |
| Map resolution (Å) | 2.83 |
| Map sharpening B factor (Å2) | -91 |
| FSC threshold | 0.143 |
| Phase randomised beyond (Å) | 3.8 |
| **Refinement** | |
| Initial model | generated from talos data |
| Protein Residues | 7,336 |
| Non-hydrogen atoms | 53,970 |
| Chains | 14 |
| Initial fitiing coordinates (PDB ID) | 2nwc |
| MolProbity score | 1.52 |
| Clashscore | 10.02 |
| Rotamer Outliers (%) | 0 |
| **Ramachandran plot** | |
| Favored (%) | 99.06 |
| Allowed (%) | 0.94 |
| Outliers (%) | 0 |

**Supplementary Table 2. DOR-GroEL-UGT1A (2.7 Å) data processing and refinement.**

| **Data processing** | **2.7Å DOR-GroEL-UGT1A** |
|---|---|
| Number of micrographs | 2951 |
| Number of frames per-micrograph | 50 |
| Applied dose per frame (e−/Å2) | 0.798 |
| Final particle images | 1,018,943 |
| Symmetry imposed | |
| Map resolution (Å) | 2.74 |
| Map sharpening B factor (Å2) | -76.71 |
| FSC threshold | 0.143 |
| Phase randomised beyond (Å) | 3.8 |
| **Refinement** | |
| Initial model | 2.8Å Consensus map |
| Protein Residues | 7,336 |
| Non-hydrogen atoms | 53,970 |
| Chains | 14 |
| Initial fitiing coordinates (PDB ID) | 2.8 Å Consensus model |
| MolProbity score | 1.54 |
| Clashscore | 10.54 |
| Rotamer Outliers (%) | 0 |
| **Ramachandran plot** | |
| Favored (%) | 98.86 |
| Allowed (%) | 1.14 |
| Outliers (%) | 0 |

**Supplementary Table 3. Structural data processing and refinement of the 3.5 Å and 3.26 Å SER-GroEL-UGT1A complexes.**

| **Data processing** | **3.5Å SER(1)-GroEL-UGT1A** | **3.26Å SER(2)-GroEL-UGT1A** |
|---|---|---|
| Number of micrographs | 2951 | 2951 |
| Number if frames per-micrograph | 50 | 50 |
| Applied dose per frame (e−/Å2) | 0.798 | 0.798 |
| Final particle images | 21,787 | 36,392 |
| Symmetry imposed | C1 | C1 |
| Map resolution (Å) | 3.52 | 3.26 |
| Map sharpening B factor (Å2) | -72.7 | -65.9 |
| FSC threshold | 0.143 | 0.143 |
| Phase randomised beyond (Å) | 8.15 | 7.9 |
| **Refinement** | | |
| Initial model | 2.8Å Consensus map | 2.8Å Consensus map |
| Protein Residues | | 7,336 |
| Non-hydrogen atoms | | 53,970 |
| Chains | 14 | 14 |
| Initial fitiing coordinates (PDB ID) | none | 2.8 Å Consensus model |
| MolProbity score | none | 1.55 |
| Clashscore | none | 10.88 |
| Rotamer Outliers (%) | none | 0 |
| **Ramachandran plot** | | |
| Favored (%) | none | 99.45 |
| Allowed (%) | none | 0.55 |
| Outliers (%) | none | 0 |

**Supplementary Table 4. OR1–4 focus map data processing and refinement.**

| **Refinement** | **OR1** | **OR2** |
| --- | --- | --- |
| Initial model | 2.7Å DOR-GroEL-UGT1A map | 2.7Å DOR-GroEL-UGT1A map |
| Protein Residues | 550 | 550 |
| Non-hydrogen atoms | 3984 | 3984 |
| Chains | | 1 |
| Initial fiting coordinates (PDB ID) | Chain K of 2.7Å DOR-GroEL-UGT1A | Chain K of 2.7Å DOR-GroEL-UGT1A |
| MolProbity score | 1.45 | 1.46 |
| Clashscore | 8.3 | 8.43 |
| Rotamer Outliers (%) | 0 | 0 |
| **Ramachandran plot** | | |
| Favored (%) | 98.85 | 99.04 |
| Allowed (%) | 1.15 | 0.96 |
| Outliers (%) | 0 | 0 |

| **Refinement** | **OR3** | **OR4** |
| --- | --- | --- |
| Initial model | 2.7Å DOR-GroEL-UGT1A map | 2.7Å DOR-GroEL-UGT1A map |
| Protein Residues | 550 | 550 |
| Non-hydrogen atoms | 3984 | 3984 |
| Chains | | |
| Initial fiting coordinates (PDB ID) | Chain K of 2.7Å DOR-GroEL-UGT1A | Chain K of 2.7Å DOR-GroEL-UGT1A |
| MolProbity score | 1.51 | 1.55 |
| Clashscore | 9.71 | 10.73 |
| Rotamer Outliers (%) | 0 | 0 |
| **Ramachandran plot** | | |
| Favored (%) | 98.66 | 98.47 |
| Allowed (%) | 1.34 | 1.53 |
| Outliers (%) | 0 | 0 |

**Supplementary Table 5. ER-1 and ER-2 focus map data processing and refinement.**

| Refinement | ER1 | ER2 |
|---|---|---|
| Initial model | 3.26Å SER(2)-GroEL-UGT1A map | 3.26Å SER(2)-GroEL-UGT1A map |
| Protein Residues | 550 | 550 |
| Non-hydrogen atoms | 3984 | 3984 |
| Chains | | |
| Initial fitiing coordinates (PDB ID) | Chain B of 3.26Å SER(2)-GroEL-UGT1A | Chain B of 3.26Å SER(2)-GroEL-UGT1A |
| MolProbity score | 1.5 | 1.43 |
| Clashscore | 9.45 | 7.79 |
| Rotamer Outliers (%) | 0 | 0 |
| **Ramachandran plot** | | |
| Favored (%) | 98.47 | 98.85 |
| Allowed (%) | 1.53 | 1.15 |
| Outliers (%) | 0 | 0 |

**Supplementary Table 6. ATP docking energies filtered by an RMSD ≤ 4.5 Å and the top three best energy-filtered RMSD ≤ 4.5 Å.**

| ER1 | | ER2 | |
|---|---|---|---|
| RMSD ≤ 4.5 | Docking Score (kcal/mol) | RMSD ≤ 4.5 | Docking Score (kcal/mol) |
| 3.69923149 | -4.563 | 4.585259864 | -4.339 |
| 2.050067701 | -4.374 | 4.16142791 | -4.182 |
| 4.200170679 | -4.11 | 4.518244513 | -4.03 |
| 4.303793109 | -3.306 | 3.769600458 | -3.252 |
| 3.468225207 | -3.282 | 4.363801833 | -2.953 |
| 4.438499728 | -3.275 | | |
| 4.474371269 | -3.197 | | |
| 3.035609791 | -2.925 | | |
| 4.300617094 | -2.01 | | |

| OR1 | | OR2 | |
|---|---|---|---|
| RMSD ≤ 4.5 | Docking Score (kcal/mol) | RMSD ≤ 4.5 | Docking Score (kcal/mol) |
| 3.969147241 | -6.215 | 3.151581173 | -6.013 |
| 4.144326364 | -5.808 | 4.04120018 | -5.912 |
| 3.778760666 | -5.681 | 3.99333072 | -5.439 |
| 4.028108572 | -5.552 | 3.829634541 | -4.781 |
| 3.556463931 | -5.455 | 4.387441423 | -4.744 |
| 3.656915302 | -5.354 | 3.611447224 | -4.667 |
| 3.717675565 | -5.327 | 3.586147355 | -4.63 |
| 3.745441189 | -5.327 | 3.552604777 | -4.575 |
| 3.689221331 | -5.226 | 4.296406666 | -4.564 |
| 4.131569206 | -5.125 | 3.458401926 | -4.445 |
| 4.13130495 | -5.074 | 4.227691436 | -4.445 |
| 3.703589004 | -5.072 | 3.319528478 | -4.401 |
| 3.639872483 | -5.009 | 3.799532992 | -4.398 |
| 4.273796466 | -4.794 | 2.532296668 | -4.384 |
| 4.223885841 | -4.687 | 3.470941936 | -4.353 |
| 4.407734854 | -4.612 | 3.701762096 | -4.182 |
| 3.901153414 | -4.611 | 3.199717556 | -4.171 |
| 3.943646027 | -4.585 | 4.147292992 | -4.168 |
| 3.836397915 | -4.578 | 3.954246723 | -4.153 |
| 3.895708856 | -4.551 | 3.436171467 | -4.132 |
| 4.021745233 | -4.546 | 4.339294839 | -3.638 |
| 3.635453692 | -4.474 | 3.895376048 | -3.188 |
| 3.551667077 | -4.347 | | |
| 4.22860023 | -3.703 | | |
| 4.262721928 | -3.146 | | |

| OR3 | | OR4 | |
|---|---|---|---|
| RMSD ≤ 4.5 | Docking Score (kcal/mol) | RMSD ≤ 4.5 | Docking Score (kcal/mol) |
| 4.263809508 | -5.826 | 4.483376867 | -5.499 |
| 4.312986399 | -5.465 | 3.784887742 | -4.917 |
| 4.387422623 | -5.225 | 3.837033602 | -4.476 |
| 3.922847185 | -5.023 | 4.453240298 | -4.369 |
| 4.196827057 | -5.018 | 4.333225667 | -4.35 |
| 3.368990978 | -4.828 | 3.721723155 | -4.029 |
| 3.59254593 | -4.579 | | |
| 3.536884486 | -4.524 | | |
| 3.32967258 | -4.343 | | |
| 4.09786385 | -4.29 | | |
| 4.475838003 | -4.285 | | |
| 3.615249446 | -3.575 | | |
| 3.487750612 | -3.31 | | |
| 3.542462181 | -2.317 | | |

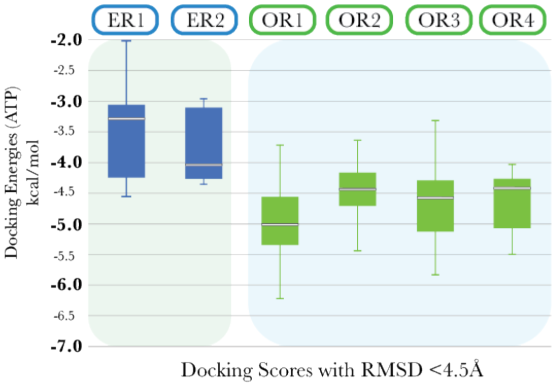

| Name | Docking Score (kcal/mol) |
|---|---|
| ER1 | -4.563 |
| ER1 | -4.374 |
| ER1 | -4.11 |
| ER2 | -4.339 |
| ER2 | -4.182 |
| ER2 | -4.03 |
| OR1 | -6.215 |
| OR1 | -5.808 |
| OR1 | -5.681 |
| OR2 | -6.013 |
| OR2 | -5.912 |
| OR2 | -5.439 |
| OR3 | -5.826 |
| OR3 | -5.465 |
| OR3 | -5.225 |
| OR4 | -5.499 |
| OR4 | -4.917 |
| OR4 | -4.476 |

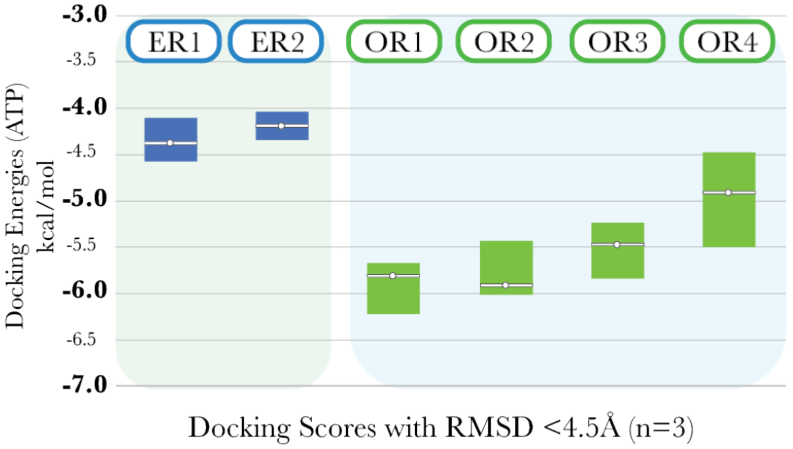
